# Looking forward does not mean forgetting about the past: ERP evidence for the interplay of predictive coding and interference during language processing

**DOI:** 10.1101/567560

**Authors:** Pia Schoknecht, Dietmar Roehm, Matthias Schlesewsky, Ina Bornkessel-Schlesewsky

**Author notes:** Corresponding author: Pia Schoknecht, Erzabt-Klotz-Straße 1, 5020 Salzburg, Austria.

## Abstract

Interference and prediction have independently been identified as crucial influencing factors during language processing. However, their interaction remains severely underinvestigated. Furthermore, the neurobiological basis of cue-based retrieval and retrieval interference during language processing remains insufficiently understood. Here, we present an ERP experiment that systematically examined the interaction of interference and prediction during language processing. We used the neurobiologically well-established predictive coding framework and insights regarding the neuronal mechanisms of memory for the theoretical framing of our study. German sentence pairs were presented word-by-word, with an article in the second sentence constituting the critical word. We analyzed mean single trial EEG activity in the N400 time window and found an interaction between interference and prediction (measured by cloze probability). Under high predictability, no interference effects were observable. Under the predictive coding account, highly predictable input is totally explained by top-down activity. Therefore the input induces no retrieval operations which could be influenced by interference. In contrast, under low predictability, conditions with high interference or with a close, low-interference distractor showed a broadly distributed negativity compared to conditions with a distant, low-interference distractor. We interpret this result as showing that when unpredicted input induces model updating, this may elicit memory retrieval including the evaluation of distractor items, thus leading to interference effects. We conclude that interference should be included in predictive coding-based accounts of language because prediction errors can trigger retrieval operations and, therefore, induce interference.

## 1. Introduction

Interference and prediction have independently been identified as crucial influencing factors during language processing. However, their interaction remains severely underinvestigated. Furthermore, despite numerous behavioral studies investigating interference during sentence processing, its neurobiological basis remains insufficiently understood. We addressed these issues with the present study: We aimed to deepen the neurobiological understanding of interference during language comprehension by studying the interplay of interference and prediction within the neurobiologically well-established predictive coding framework. In the remainder of the introduction, we introduce interference during language processing and the sources of neurobiological grounding of our study: Neuronal mechanisms of memory representations and the predictive coding framework including the neurobiology of the N400 component. Then we turn to the interplay of interference and prediction and finally to the present study.

### 1.1 Interference during Language Processing

To successfully process a sentence or discourse, it is necessary to access previously encountered words, so that new words can be integrated with ones that have already been processed. The cognitive mechanisms underlying this process have been modeled in detail within the cue-based parsing framework (Lewis, Vasishth, & Van Dyke, 2006; Van Dyke & Lewis, 2003). (Please note that we use the cue-based parsing models put forward by Lewis and Vasishth (2005) and McElree (2000) interchangeably. The two models only differ in aspects of retrieval latency which are not essential to the presented study; (see Nicenboim & Vasishth, 2018 for detailed discussion of the models). The cue-based parsing framework views memory retrieval as a cue-based process, which compares retrieval cues generated at the retrieval site with the features of the words in memory. Each new word or constituent is encoded into memory as a feature bundle (e.g. case, number, gender etc.). Importantly, not only the constituent itself is represented, but also predicted constituents that are required to form a grammatical sentence. Thus, when the noun *toy* is encountered in example 1 (from Lewis et al., 2006), two memory representations are constructed: one depicting its features and the other a prediction of the corresponding predicate (verb). When the verb *arrived* is processed, retrieval cues are generated to integrate it into the sentence context. The cues are matched in parallel against all (recent) memory representations. A sufficient match results in retrieval of the predicate prediction and the integration of verb and noun.

(1) Melissa knew that the toy from her uncle in Bogotá arrived today.

But retrieval and, hence, comprehension difficulty can arise due to similarity-based interference from cue overlap, i.e. an overlap between the retrieval cues and features of more than one word / prediction recently stored in memory (Lewis et al., 2006). The correct to-be-retrieved word / prediction is called the target of the retrieval and other words / predictions in memory may function as interfering distractors during the retrieval process. Numerous behavioral studies have provided evidence of similarity-based interference (see Jäger, Engelmann, and Vasishth (2017) for a recent review and meta-analysis). For example, Van Dyke and McElree (2006) used a dual-task paradigm to study interference from extra-sentential memory items (see 2a) during sentence comprehension. They presented it-clefts (see 2b) that required a retrieval operation to integrate the matrix sentence verb (*sailed* / *fixed*) with its object (*boat*). The object is the target of the retrieval.

(2a) Memory load set: table – sink – truck

(2b) It was the boat that the guy who lived by the sea <u>sailed</u> / <u>fixed</u> in two sunny days.

In load conditions, three nouns functioning as distractors (cf. 2a) were presented before a self-paced sentence reading task (for sentences such as 2b). In high interference conditions, the distractors were plausible objects of the matrix verb, and in low interference conditions, they were implausible (i.e. it is possible to fix, but not to sail a table / sink / truck). To ensure that there were no confounding differences in the comprehension of the verbs, no load conditions, in which participants just read the sentences without an additional task, were also included. Reading times for the critical verb carrying the interference manipulation were longest for high interference conditions with memory load. These results were interpreted as showing that retrieval is the locus of interference during sentence comprehension rather than encoding, as the retrieval context was manipulated while encoding was held constant (Van Dyke & McElree, 2006).

Van Dyke (2007) showed that distractors that match only a subset of retrieval cues could nevertheless influence retrieval and cause processing difficulties. Furthermore, it was found that distractors occurring between target and retrieval site (retroactive interference) have more impact on processing than distractors that are processed before the target (proactive interference) (Martin & McElree, 2009).

While cue-based retrieval and interference in sentence processing are well established from a behavioral perspective, their neurobiological basis remains insufficiently understood. Initial ERP evidence was presented by Martin, Nieuwland, and Carreiras (2012), who studied how distractors affect the processing of NP ellipses in Spanish (see 3). To successfully process the ellipsis, the antecedent *camiseta* (‘t-shirt’) needs to be retrieved at the position of the determiner *otra* (‘another’). The authors manipulated the (mis)match between the gender of the target (*camiseta*, ‘t-shirt’ [feminine]), the distractor (*falda*, ‘skirt’ [feminine] / *vestido*, ‘dress’ [masculine]) and the retrieval cues generated at the retrieval site (*otra*, ‘another’ [feminine] / *otro*, ‘another’ [masculine]).

(3) Marta se compró la camiseta que estaba al lado de la falda / del vestido y Miren cogió otra / otro […] para salir de fiesta.

‘Marta bought the t-shirt [feminine] that was next to the skirt [feminine] / dress [masculine] and Miren took another [feminine] / another [masculine] to go to the party.’

They reported a sustained, broadly distributed negativity for ungrammatical ellipses (*camiseta* [feminine] … *otro* [masculine]) compared to grammatical ellipses (*camiseta* [feminine] … *otra* [feminine]). Within grammatical ellipses, this effect was modulated by the gender of the distractor: a mismatching distractor (*vestido*, ‘dress’ [masculine]) led to a negativity compared to matching distractor (*falda*, ‘skirt’ [feminine]). It was concluded that the processing of grammatical ellipses, i.e. the retrieval of the antecedent / target, was interrupted by the mismatching features of a mismatching distractor. In a follow-up study, Martin, Nieuwland, and Carreiras (2014) changed the syntactic structure of their material to render the antecedents of the ellipses (the target of the retrieval) less distinct. In this study, grammatical ellipses with matching distractors elicited an early anterior negativity compared to grammatical ellipses with mismatching distractors and ungrammatical ellipses. This was regarded as an effect of similarity-based interference. Thus, while both studies found negativities, they were elicited by different conditions (matching vs. mismatching distractors), had different distributions and latencies. The neurocognitive correlates of interference therefore clearly require further investigation. In addition, no attempt has been made to date to elucidate the possible neurobiological mechanisms underlying retrieval interference during language comprehension.

### 1.2. Neurobiological Grounding: Neuronal Memory Representations, Predictive Coding and the N400

With the present study, we aimed to deepen the neurobiological understanding of interference during language comprehension in two ways: a) by including considerations regarding the neuronal basis of memory (Jonides et al., 2008) and b) by studying the interplay of interference and prediction within the neurobiologically well-established predictive coding framework (Friston, 2005).

Despite the above-mentioned lack of neurobiological grounding of interference during language processing, Jonides et al. (2008) provided insights into the neuronal processes underlying short-term memory and consequently similarity-based interference as a cause of forgetting. Items that are available for cognitive processing without retrieval processes are in the attentional focus (see McElree (2006) for a detailed discussion of the capacity limit of the attentional focus); these focused items are represented by a synchronized firing pattern of neurons in primary and secondary association cortex that equal the activation during perception or action. “In short, item representations are stored where they are processed.” (Jonides et al., 2008, p. 213). Items in memory outside the focus are supported by short-term synaptic plasticity in the corresponding areas. Therefore, primarly posterior brain regions are active during storage intervals in memory tasks, while frontal regions undertake executive functions such as attention shifts, retrieval and resolution from interference.

Jonides et al. (2008) offer neuronal explanations for both phenomena that have been broadly discussed as causes of forgetting: interference and decay. As representations consist of temporally synchronized neuronal firing, decay may be explained by variability in the firing rates of single neurons which subsequently fall out of synchrony, rendering the representation firing pattern less coherent and therefore, difficult to discriminate from noise. When the noise is not random, but has been structured by previously encountered items which are similar to the current item, the noisy firing pattern and the firing pattern making up the item representation overlap and compete (proactive interference). This results in an increased likelihood of the corresponding neurons falling out of synchrony or in a less coherent firing pattern of the item representation in the first place, which falls out of synchrony more quickly. Retroactive interference is induced by the firing pattern of a new item disturbing the synaptic weights of the previous item based on overlap. Thus, similarity-based interference – both pro- and retroactive – is caused by structured noise in the neural pattern. Here it is important to note that retroactive interference has been found to be more severe during language comprehension than proactive interference (Martin & McElree, 2009).

This account views retrieval as a process mediated by frontal control which corresponds to the reactivation of the target’s neuronal firing pattern in posterior areas and medial-temporal areas which support contextual binding. The reactivation of the firing pattern might be carried out by emergent pattern-completion of attractor networks (Hopfield, 1982) which is complicated by the above-mentioned causes of noise (Jonides et al., 2008). This view can be applied to interference paradigms in language processing (see *1.1 Interference during Language Processing*) by assuming that the retrieval cues generated at the retrieval site activate the corresponding parts of the firing pattern (e.g. gender, number or semantic features like being sailable) of the target item representation. This partial activation would reinstantiate the firing pattern of the previously focused target item representation which was still representated in synaptic weights. Additionally, we assume that the prediction of an item also leads to an attentional shift towards that item and thereby to the instantiation of its firing pattern as if that item would be perceived at the moment of prediction. For a schematic illustration of the described neuronal processes, see Figure 1.

**Figure 1:**
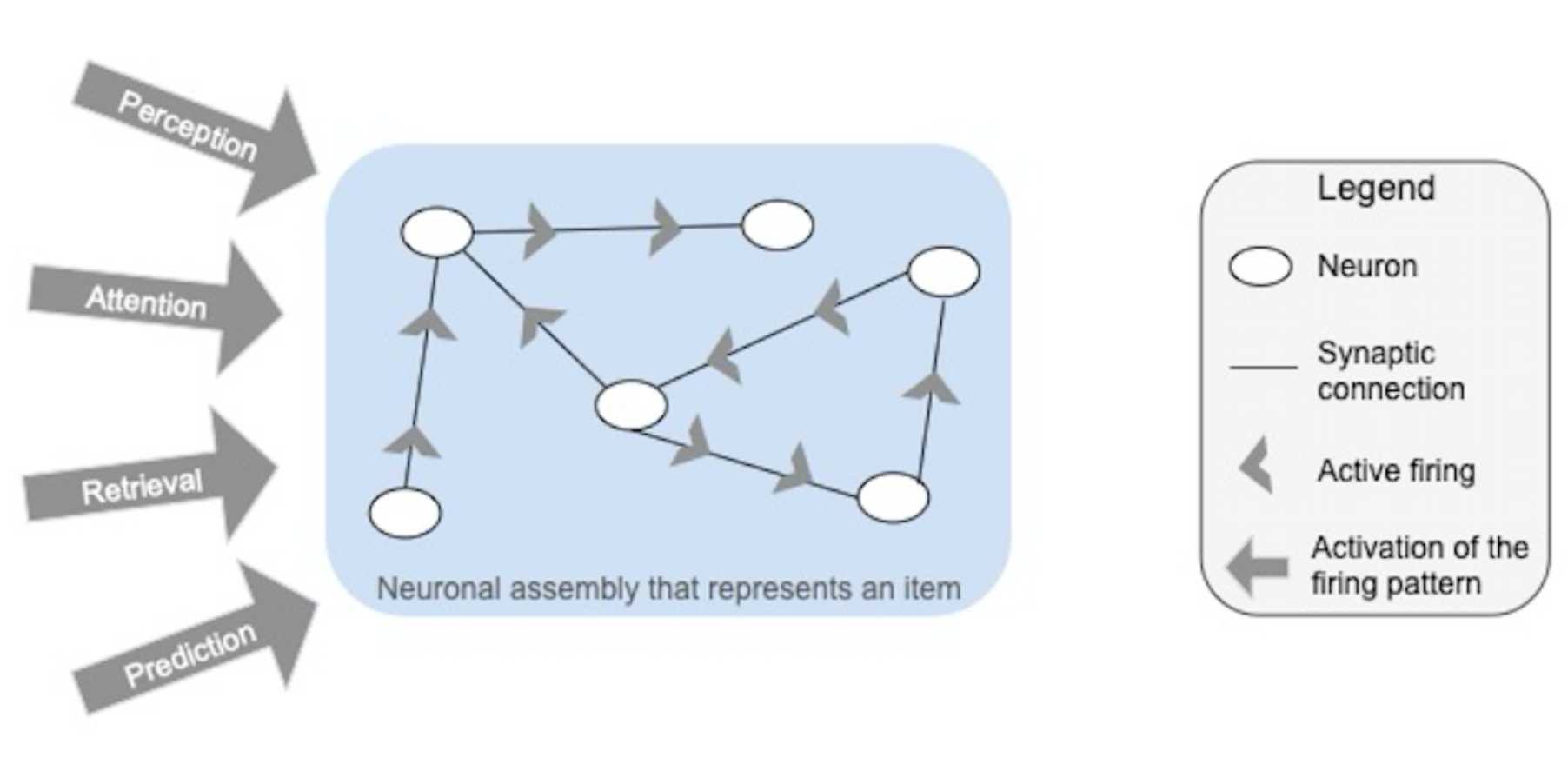
Schematic illustration of a neuronal assembly making up the representation of an item in memory. As long as the item is not in the attentional focus, representation is carried by synaptic connections between corresponding neurons. Perception, attention, retrieval or prediction of / to the item equally lead to active firing of the neurons within the assembly, whereby the item enters the attentional focus.

Now we turn to our second source of neurobiological grounding: prediction within the neurobiologically well-established predictive coding framework. Predictive coding states that prediction is a general principle of brain function, with generative predictive models serving to explain the causes of sensory input (Friston, 2005). If true, this means that the brain also engages in predictive modeling of all linguistic input (e.g. Bornkessel-Schlesewsky, Schlesewsky, Small, & Rauschecker, 2015; Freunberger & Roehm, 2017; Kandylaki et al., 2016; Lewis & Bastiaansen, 2015; Pickering & Garrod, 2007). Predictive processing is assumed to be organized within a hierarchically structured cortical architecture, within which anatomical feedback connections (from higher to lower levels) communicate predictions. Feedforward connections (from lower to higher levels), by contrast, communicate sensory input or – more precisely – only the parts of the input that are divergent from the prediction (Clark, 2013; Rao & Ballard, 1999). Thus, prediction errors essentially serve as a proxy for sensory information (e.g. Feldman & Friston, 2010), since the feed-forward activity associated with sensory input is “silenced” when predicted (i.e. excitatory bottom-up activity is countered by inhibitory top-down activity in this case; cf. Friston, 2010). Overall, the system strives to reduce prediction errors, but when they do occur, they are used to update the current model so that more accurate predictions can be generated in the future.

According to a recent proposal by Bornkessel-Schlesewsky and Schlesewsky (2019), the N400 event-related potential provides a window on predictive coding during language comprehension. Specifically, this account posits that the N400 indexes *precision-weighted* prediction errors, with precision defined as the inverse of variance (Feldman & Friston, 2010) and – in the context of language – equating to the relevance of a particular information source for the form-to-meaning mapping. A prediction has high precision when the information source from which it is derived has high certainty, i.e. the information source provides a direct link between form and meaning. In contrast, low precision predictions stem from information sources with (high) uncertainty. Higher precision prediction errors are associated with more pronounced N400 amplitudes (cf. Todd et al., 2014, for discussion of a similar precision-weighted mechanism in the context of the mismatch negativity, MMN).

This account offers an explanation for the presence versus absence of N400 effects for the same, prediction-error inducing phenomenon across different languages, depending on the language specific relevance (or “validity”, cf. MacWhinney, Bates, & Kliegl, 1984) of cues such as word order, animacy or case marking (see Bornkessel-Schlesewsky et al. (2011) for empirical evidence, and Bornkessel-Schlesewsky and Schlesewsky (2019) for detailed discussion). In summary, while N400 amplitude shows a general tendency to decrease for predicted stimuli and increase for stimuli that involve prediction error (see e.g. Dröge, Fleischer, Schlesewsky, & Bornkessel-Schlesewsky, 2016; Frank, Otten, Galli, & Vigliocco, 2015; Kuperberg, 2016), the magnitude of the N400 effect is assumed to be proportional to the degree to which the prediction error actually leads to a model update. In the present study, we will draw on this proposal to formulate our hypotheses with regard to the relation between prediction and interference during language processing.

### 1.3. Interplay of Prediction and Interference

The interplay of prediction and interference has — to the best of our knowledge — only been examined by one study to date. Campanelli, Van Dyke, and Marton (2018) used the Van Dyke and McElree (2006) paradigm (cf. example 2, above) with an added predictability manipulation (see example 4). By varying the subject noun (*person* / *choreographer*), the authors manipulated the cloze probability (CP), and thereby the predictability, of the critical matrix sentence verb.

(4a) Memory load set: website – handbag – password

(4b) It was the dance that the person / choreographer who lived in the city <u>performed</u> / <u>created</u> early last month.

Campanelli et al. (2018) observed (marginal) reading time differences related to predictability and interference only in the sentence-final spillover region (e.g. *early last month*). In the load conditions, residual log reading times were fastest in interference conditions with high predictability and slowest in interference conditions with low predictability of the critical verb. This result provides an initial indication that interference effects are canceled out when the word at the retrieval site is highly predictable. However, Campanelli et al. (2018)’s design was not fully crossed, as they investigated two levels of prediction (high, low) within the interference conditions but only low prediction within no interference conditions. Thus, it is not possible to fully characterize the interplay between prediction and interference on the basis of these results.

We can, nevertheless, put forward a hypothesis regarding the relationship between prediction and interference within the context of a predictive coding framework. Recall from our discussion of predictive coding above that predictions essentially correspond to top-down activity within a hierarchically organized processing architecture, and that this activity cancels out bottom-up, stimulus-related activity when all features of a stimulus are fully predictable – thereby preventing the occurrence of prediction errors. By contrast, interference arises when bottom-up information calls for a retrieval and the features of the retrieval target are too similar to those of other items currently held in memory. It thus appears plausible that retrieval operations are only required in the absence of perfect predictability, i.e. when it is not possible to fully cancel out stimulus-related activity. In other words, when an upcoming word is fully predicted (and therefore its neuronal firing pattern was already totally activated before it was encountered), it does not contain any new information that could serve to initiate a retrieval operation. We sought to examine this hypothesis with the present study.

### 1.4. Present Study

The present study examined the interplay between prediction and interference in German, using a design that fully crossed the two factors. We focused on German articles as critical stimuli within a broader (two) sentence context, because they overtly mark case, number and gender and are thus ideal for manipulating interference. Despite the design opportunities afforded by German for the examination of interference, to the best of our knowledge, only one study has investigated retrieval interference during German sentence processing (Nicenboim, Vasishth, Engelmann, & Suckow, 2018). The authors used a self-paced reading paradigm and found weak evidence for an interference effect during the retrieval of a subject in the presence versus absence of distractor nouns that shared the same number feature. Here, we aimed to provide further evidence for interference effects in German in addition to presenting what is – to the best of our knowledge – the first ERP study to investigate the interplay of predictive coding and interference. Specifically, we manipulated gender interference and prediction during the processing of an article (see example 5). In German, this word class carries (only) case, number and gender information.

(5) Im Glas liegt eine Raupe und in der Schachtel sitzt ein Käfer. Peter befreit <u>den</u> Käfer aus der Schachtel.

‘In the glas there lies a caterpillar [feminine] and in the box there sits a beetle [masculine]. Peter frees the [masculine] beetle from the box.’

The target (*Käfer* [masculine], ‘beetle’) and a distractor (*Raupe* [feminine], ‘caterpillar’) were introduced in a context sentence. At the critical article position within the second (target) sentence (*den* [masculine], ‘the’), the language processing system should seek to establish coreference with a noun in the preceding context (see e.g. Schumacher, 2009). The article therefore constitutes the retrieval site of interest. We hypothesized that the retrieval process would be influenced by both the predictability of the article and that of the following noun, as well as by the degree of interference from the distractor noun. We decided to also include the predictability of the following noun as a predictor variable, because we assumed that some processes taking place during the processing of the article would be affected by the predictability of the noun. These include, for example, the generation of a noun prediction or, if a prediction for the noun existed before the position of the article, whether this prediction is further supported or falsified by the article via its (in)compatibility to the predicted noun. Any or all of these processes might substantially affect the processing of the article. Materials were constructed so that they varied in terms of both target article and noun cloze probability and the degree of interference (see Methods for further details). By extending the distance between the retrieval site and the to-be-retrieved word across a sentence boundary, we hoped to gain insights into the scope of potentially interfering distractors.

### 1.5. Hypotheses

Our hypotheses were as follows: when the article is fully predictable, prediction confirmation at this position further supports the existing prediction of the following noun. In cases of prediction disconfirmation at the article, by contrast, the internal model needs to be adjusted to prevent further prediction errors and the prediction for the upcoming noun needs to be revised. This revision necessarily involves memory retrieval of potential candidates and we hypothesized that it should be influenced by interference. Following Bornkessel-Schlesewsky and Schlesewsky (2019), we expected to find N400 effects for prediction errors versus prediction confirmation at the article and hypothesized that these would be modulated by interference. Interference was assumed to shape model updating as it influences the precision of the gender feature of the article. In low interference conditions, the gender feature unambiguously matched one NP in the context and, therefore, was a cue with high precision. In contrast, in high interference conditions, the gender feature of the article rendered it ambiguous as it matched the target and distractor NP, so that in these conditions the gender feature was associated with low precision.

## 2. Materials and Methods

### 2.1. Participants

Forty-four right-handed participants (8 male, mean age: 22.75 years, range: 18 – 35) gave written informed consent and received either course credit or 20 Euros for their participation. All were native (Austrian) German speakers with normal or corrected-to-normal vision and no history of neurological or psychiatric disorders. None of the participants took part in the questionnaire pretest. Four participants had to be excluded because of technical issues, three were excluded because their response accuracy was below 75 % and five participants were excluded because of excessive artifacts (more than 25 % of the critical sentences were affected). The data of thirty-two participants (5 male, mean age: 22.34 years, range: 18 – 35) was used for final analyses.

### 2.2. Materials

Eighty-five sets of four sentence pairs were constructed (see Table 1 for an example). Two noun phrases (NPs), which either matched or mismatched according to their gender features, were introduced in a context sentence and one of them was repeated in the target sentence. Target sentences were identical across conditions. The repeated NP was always the object of the target sentence and its article was our critical position for ERP analyses. In German, articles and corresponding nouns must have congruent gender, number and case features. In the high interference conditions (1 / 2), the article could grammatically be followed by each of the two nouns in the context, while, in the low interference conditions (3 / 4), there is only one compatible noun in the context. Additionally, the distance between the target noun and the retrieval site (the article) was manipulated via the word order in the context sentence; resulting in retroactive interference (target … distractor … retrieval site) in the long target distance conditions and proactive interference (distractor … target … retrieval site) in the short target distance conditions.

**Table 1:**
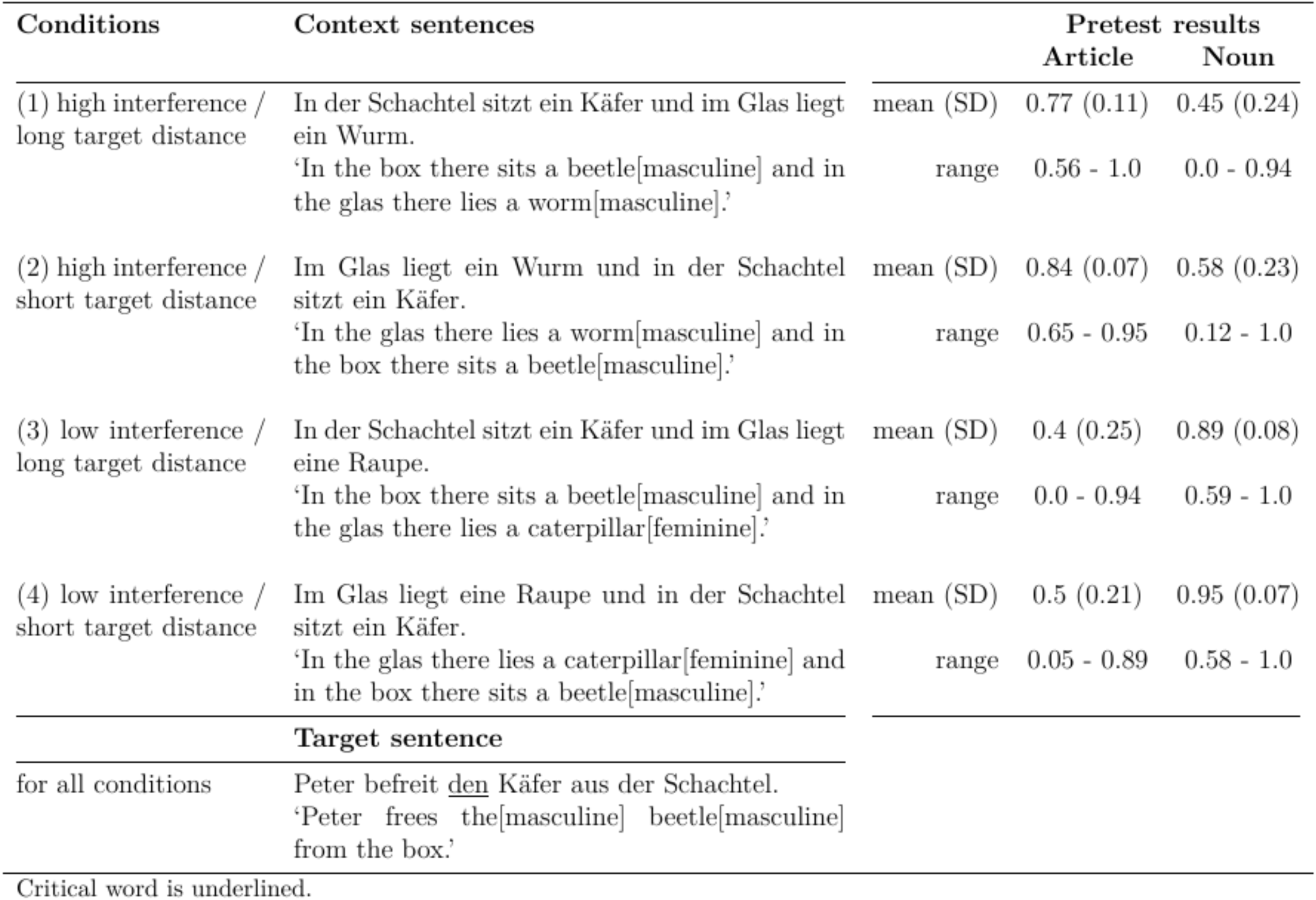
Example material and pretest results (cloze probability values) for the critical article and noun within the target sentence displayed beside the corresponding context sentences.

### 2.3. Questionnaire Pretest

A sentence completion pretest was conducted to obtain cloze values for the critical article and the following noun in context (Kutas & Hillyard, 1984; Taylor, 1953). The material was split into four lists, with only one sentence pair from each lexical set occurring per list. In addition, there were two versions of each list: one truncated the target sentence before and one after the critical article, resulting in eight different questionnaires. The different lists / versions were constructed to ensure that the pretest participants stayed naïve towards the manipulation. No fillers were included in the pretest. A total of 144 university students participated in the pretest (16 to 20 per questionnaire version; 39 male, mean age = 22.5 years, range: 18 – 40). Pretest participants were native speakers of (Austrian) German and were instructed to intuitively complete the sentence pairs. None of the pretest participants also took part in the main study.

The obtained cloze probability (CP) values ranged from 0 to 1 across conditions for both the article and noun. The results of the pretest are shown in Table 1. Pairwise contrasts were computed with Welch’s t-test in `R` (R Core Team, 2019, Version 3.6.0). The CP of the article in the high interference conditions was higher overall than in the low interference conditions (long target distance: diff = 0.37, p < 0.001, 95 % CI = 0.31 – 0.44; short target distance: diff = 0.34, p < 0.001, 95 % CI = 0.28 – 0.38). This is a logical consequence of the fact that, no matter with which of the two nouns of the high interference context the participants chose to continue the sentence, the article had the same form because the nouns shared the same gender feature. In contrast, the CP values for the article in the low interference conditions were lower because the article form varied based on the decision regarding which of the context nouns should follow. In both interference conditions, the participants showed a recency effect, i.e. they chose an article congruent to the last-mentioned noun more often than to the first-mentioned noun (cf. the higher CP values for a short distance to referent across conditions; high interference: diff = 0.07, p < 0.001, 95 % CI = 0.03 – 0.09; low interference: diff = 0.1, p = 0.005, 95 % CI = 0.03 – 0.18). In the high interference conditions, this recency effect is difficult to interpret as the nouns share the same gender and might be due to an implicit bias related to verb semantics, i.e. in the way that one of the nouns is closer associated with the verb.

The CP values for the target noun are lower overall in the high interference conditions compared to low interference conditions (long target distance: diff = - 0.44, p < 0.001, 95 % CI = -0.49 – -0.38; short target distance: diff = -0.37, p < 0.001, 95 % CI = -0.43 – - 0.32). This reflects the fact that the article is ambiguous in the high interference conditions and that, in the low interference conditions, the article only matches one noun. Like the article results, the noun CP results show a recency effect (high interference: diff = 0.13, p = 0.001, 95 % CI = 0.05 – 0.2; low interference: diff = 0.06, p < 0.001, 95 % CI = 0.04 – 0.09).

The distribution and combinations of the CP values, respectively, are shown in Figure 1. In Figure 1, it is apparent that our materials contain no sentence pairs in which a low cloze article was followed by a low cloze noun. A low cloze article was always followed by a high cloze noun: Therefore, although a specific article may have been unpredicted within the target sentence, when this article was processed, the following noun became highly predictable. These sentence pairs always belonged to low interference conditions. Therefore, the observed distribution pattern as visualized in Figure 1 is caused by the above-mentioned likelihoods of articles and nouns within the low interference conditions. In contrast, high cloze articles were followed by nouns with varying cloze values. The combination of high cloze articles and low cloze nouns is only seen for the high interference condition. Again, this is a consequence of the above-mentioned likelihoods of articles and nouns within the high interference condition. Only the combination of high cloze articles and high cloze nouns is apparent in all conditions, which can be explained by a clear preference for one of the NPs, regardless of condition. This is likely due to verb bias.

In summary, for both the article and the noun, we see that the participants of our pretest used the likelihood of an article and noun with a specific gender feature to complete the sentence pair. This likelihood was mostly determined by the characteristics of the experimental conditions. Also, for both the article and the noun, CPs reveal an offline recency effect. The participants expected the last-mentioned noun to be repeated in the target sentence. Despite these general tendencies, items differed from each other and we observed effects that might be due to item-specific verb biases. The pretest results were used as predictors in our linear mixed model analyses, which allow unbalanced designs (for details, see 2.6. Statistical Analysis).

### 2.4. Procedure

Eighty of the original eighty-five items were presented in the main study. The exclusion of five items was used to achieve an even number of items and was not based on pretest results. Two pseudo-randomized lists were constructed so that each participant saw only two versions of one item (either version 1 and 4 or version 2 and 3). To avoid order effects, half of the participants were presented a backward version of the lists. Each of the lists consisted of 240 sentence pairs (160 critical and 80 filler sentence pairs). The first sentences of the filler sentence pairs were identical to the structure of the critical sentence pairs. The second sentence of the fillers was constructed to be coherent to the first sentence but did not include a referential link to it. All sentence pairs (both critical and filler pairs) were grammatical. A third of all sentence pairs in both lists (80 out of 240) was followed by a “yes”/”no” comprehension question. There were three types of questions: a) about the action or state of one NP within the context sentence (*Liegt der Wurm im Glas?* ‘Is the worm in the glas?’), b) about which of the NPs was used in the target sentence (*Befreit er den Käfer?* ‘Does he free the beetle?’) or c) the content of the target sentence after the critical words (*Befreit er ihn aus der Schachtel?* ‘Does he free it from the box?’). Please note, that the provided example questions are hypothetical questions that correspond to the example item presented in Table 1. Each participant was asked to answer at most one question per item. Questions were not repeated within lists. Half of the questions required a “yes” answer indicated by a right or left mouse click (balanced across participants).

Participants were instructed to silently read the sentence pairs, which were presented on the center of a computer screen in a word-by-word manner. Words were presented in white letters on a black background and participants were asked to minimize eye movements, blinks and general movements. A trial began with a 400 ms fixation cross followed by 200 ms blank screen. Words were presented for 400 ms each with a 100 ms inter-stimulus interval between words; after the last word of the first sentence in the pair, there was a 1000 ms inter-stimulus interval. To encourage participants to read carefully, a third of the trials was followed by a “yes”/”no” comprehension question, which the participants had five seconds to answer. After that (or in trials without a task), there was a self-paced inter-trial interval, during which participants could blink as much as needed before pressing a button to start the next trial. All sessions began with a five-sentence-pair practice block to familiarize participants with the procedure. The following six experimental blocks were separated by short breaks. Each experimental session lasted approximately 2.5 hours; participants spent approximately 75 minutes on task. An ethics committee approved the procedure that was applied in the present study.

### 2.5. Electroencephalogram recording and preprocessing

The electroencephalogram (EEG) was recorded from 58 active scalp electrodes positioned according to the standard 10-20 system and attached to an elastic cap (ActiCap from Easycap GmbH, Herrsching, Germany). EEG recordings used an ActiCHamp amplifier (Brain Products GmbH, Gilching, Germany) with a sampling rate of 500 Hz and AFz as the ground electrode. Six electro-oculogram (EOG) electrodes were positioned above and below both eyes and at the outer canthus of each eye. Impedances were kept below 10 kΩ. All electrodes were online referenced to the left mastoid and re-referenced to the average of both mastoids offline. All preprocessing steps were carried out using the Brainvision Analyzer 2. The raw data were filtered with a bandpass of 0.1 – 30 Hz. Artifact rejection was carried out semi-automatically, based on Brainvision Analyzer 2’s Max-Min criterion. According to this criterion, intervals were rejected in which the six EOG channels showed a difference of more than 100 µV within 200 ms windows, plus an additional 200 ms window before and after the artifact.

ERPs were computed in epochs of -200 to 1000 ms relative to critical article onset (noun onset was 500 ms after article onset). While we present grand average ERPs (see Figure 2) for comparability to previous research, the current design required single trial-based analyses, as we outline below.

**Figure 2:**
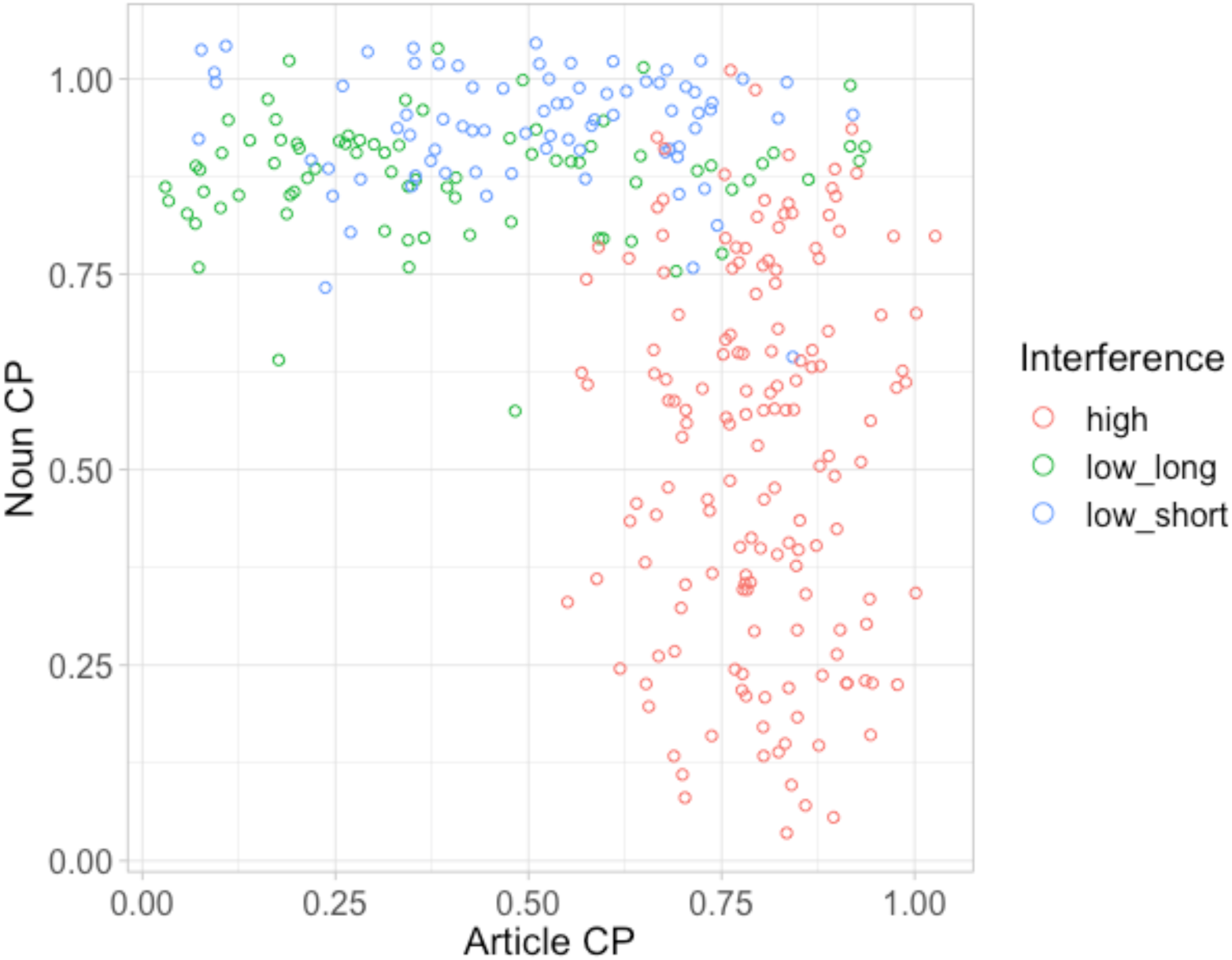
Distribution and combinations of CP values for critical article and following noun. Each circle represents one condition of one item. The number of red circles is higher because we did not differentiate between the two target distance levels for the high interference conditions.

### 2.6. Statistical Analysis

We performed linear mixed model analyses using the `R` (R Core Team, 2019, Version 3.6.0) package `lme4` (Bates, Maechler, Bolker, & Walker, 2015, Version lme4_1.1-21) on the basis of the mean activity of EEG single trial data between 300 and 500 ms post onset of the article. We chose the standard N400 time window (Kutas & Federmeier, 2011) based on theoretical considerations and our hypotheses, rather than via visual inspection of ERP waveforms. We included the topographical factors sagittality (anterior: F7/8, F5/6, F3/4, F1/2, Fz, FC5/6, FC3/4, FC1/2, FCz; central: T7/8, C5/6, C3/4, C1/2, Cz, TP7/8, CP5/6, CP3/4, CP1/2, CPz; posterior: P7/8, P5/6, P3/4, P1/2, Pz, PO7/8, PO3/4, POz) and laterality (left: F7, F5, F3, FC5, FC3, T7, C5, C3, TP7, CP5, CP3, P7, P5, P3, PO7, PO3; medial: F1/2, Fz, FC1/2, C1/2, Cz, CP1/2, CPz, P1/2, Pz, POz; right: F8, F6, F4, FC6, FC4, T8, C6, C4, TP8, CP6, CP4, P8, P6, P4, PO8, PO4). Interference was modeled as a three-level factor (high, low with short distance, low with long distance between target and retrieval site) because, in the high interference conditions, the target noun cannot be determined at the article and therefore its recency is unclear until the noun position (i.e. both conditions are identical up to the position of the article). Using the `R` package `car` (Fox & Weisberg, 2011, Version carData_3.0-2), categorical variables were encoded with sum contrasts, such that the estimate for a given fixed effects level represents the difference between this level and the grand mean (for detailed discussion of contrast coding, see Schad, Vasishth, Hohenstein, and Kliegl (2019)). Offline cloze probability values of the article and following noun were included as continuous predictors as well as the z-transformed baseline activity from 200 ms pre-stimulus until stimulus onset (Alday, 2019); this corresponds to the N400 interval of the preceding word, i.e. the target sentence verb. In addition, the model included random intercepts for both subjects and items. While the use of maximal random effects structures has been encouraged for psycholinguistic research by some authors (Barr, Levy, Scheepers, & Tily, 2013), we chose to employ random intercepts only for the present study because we focus on population-level effects rather than inter-individual differences as the investigation of the interaction between interference and prediction is novel.

For brevity, we present Type-II Wald χ^2^ tests (also provided by the `car` package) but also model summaries. We analyze effects of interest with estimated marginal means (including 95 % confidence intervals) that were computed with the `R` package `emmeans` (Lenth, 2019, Version emmeans_1.3.5.1) and visualize them with the packages `ggeffects` (Lüdecke, 2018, Version ggeffects_0.11.0) and `ggplot2` (Wickham, 2016, Version ggplot2_3.1.1). The plots also serve to resolve interactions. Prior to the analyses, we excluded two items that were inconsistently constructed, leaving us with the data for 78 items.

As discussed in section 2.3, the distribution of CP values for the article and noun obtained in the pretest does not cover all possible combinations. Accordingly, parts of the linear mixed model outputs presented in section 3 reflect estimates for certain parameter combinations for which no actual data points exist in the present study.

## 3. Results

The data and analysis scripts for this experiment can be accessed at https://osf.io/ju4mr/?view_only=4cddc81fca954fc3b99eea0f00efcef3.

### 3.1 Comprehension Question Accuracy

Participants were encouraged to answer the comprehension questions correctly and within the time limit (five seconds). As noted above, three participants were excluded for accuracy rates below 75%. For the remaining participants, comprehension accuracy was good (mean: 85.4 %, range: 76.2 – 95 %), thus demonstrating that they processed the sentences attentively. Generalized linear mixed models with one or both of our condition factors (Interference and Distance between target and retrieval site) as predictors showed no improvements compared to a base model that included only an intercept term (comparison base model to Interference model: χ^2^ = 3.01, *p* = 0.08; comparison base model to Distance model: χ^2^ = 0.01, *p* = 0.9; comparison base model to interaction of Interference and Distance model: χ^2^ = 3.9, *p* = 0.27), hence comprehension accuracy was affected neither by Interference nor by Distance. We also checked whether there was a speed accuracy tradeoff. The corresponding generalized linear mixed model (with log reaction time, Interference, Distance and their interactions as predictors) revealed that correct answers were associated with a faster response time than incorrect answers (β = -1.9, SE = 0.24, z = -7.96, p < 0.001). No other effects in this model reached significance.

### 3.2. Traditional ERP waveforms

For sake of completeness, we present traditional grand average ERP waveforms (N= 32) in Figure 2. Article onset is at 0 ms and noun onset at 500 ms. The traditional ERP methodology cannot account for the complexity of our design with continuous predictors (cf. Alday, Schlesewsky, & Bornkessel-Schlesewsky, 2017). Thus, while visual inspection of the grand average ERP waveforms is suggestive of some condition-based differences in the N400 time window at the article position, we will not interpret these further, as they do not reflect our full design.

**Figure 2:**
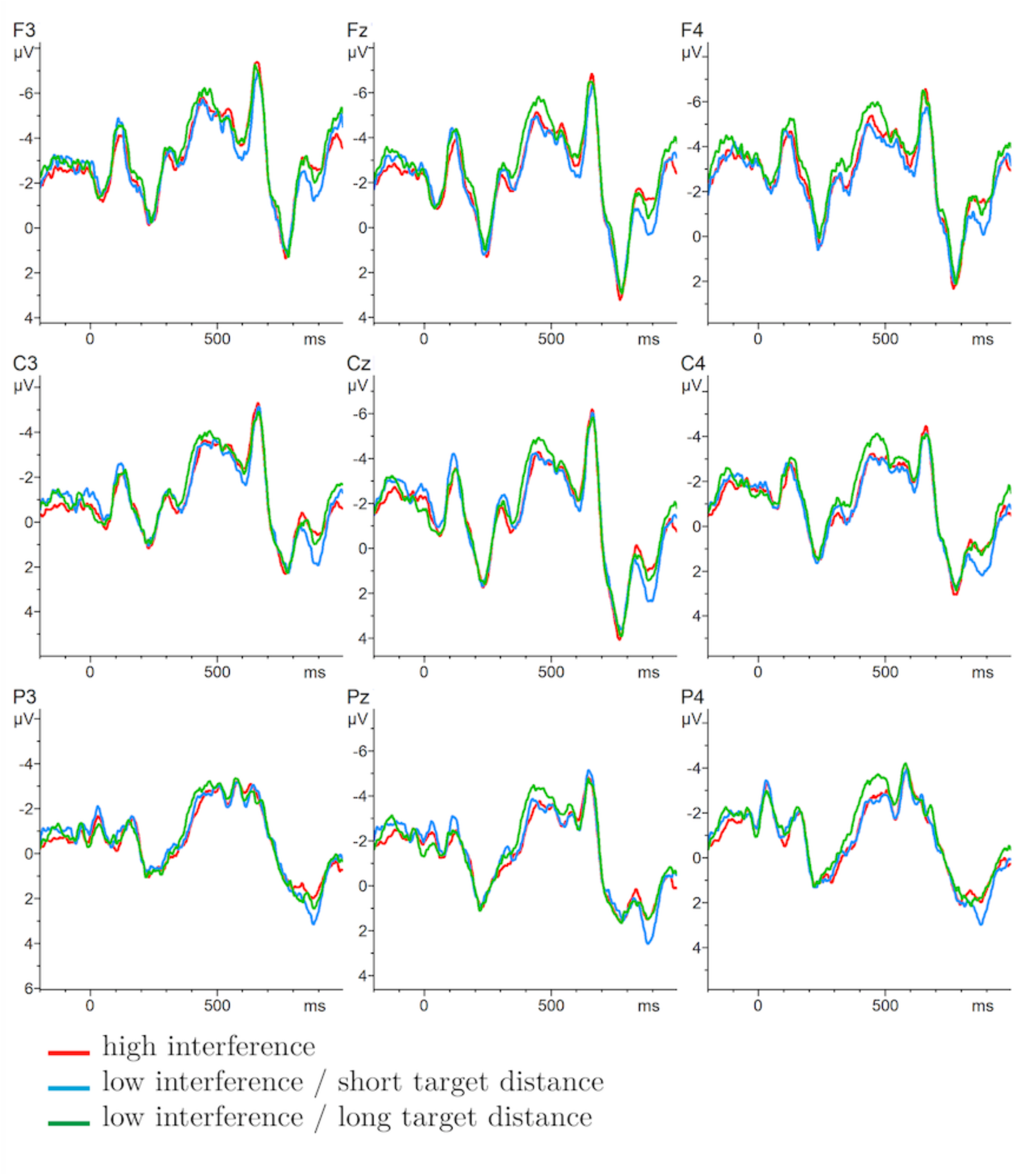
Grand averages at nine selected electrode sites. Negativity is plotted upwards.

### 3.3. Electrophysiological Data

The summary of the linear mixed model for the article is shown in Tables 2 and 3. Type-II Wald test results are shown in Table 4, we focus on the three-way interaction between Interference, Article CP and Noun CP (χ^2^(2) = 26.7, *p* < 0.001), which is visualized in Figure 3. The absence of an interaction with sagittality or laterality indicates that the effects are broadly distributed.

**Figure 3:**
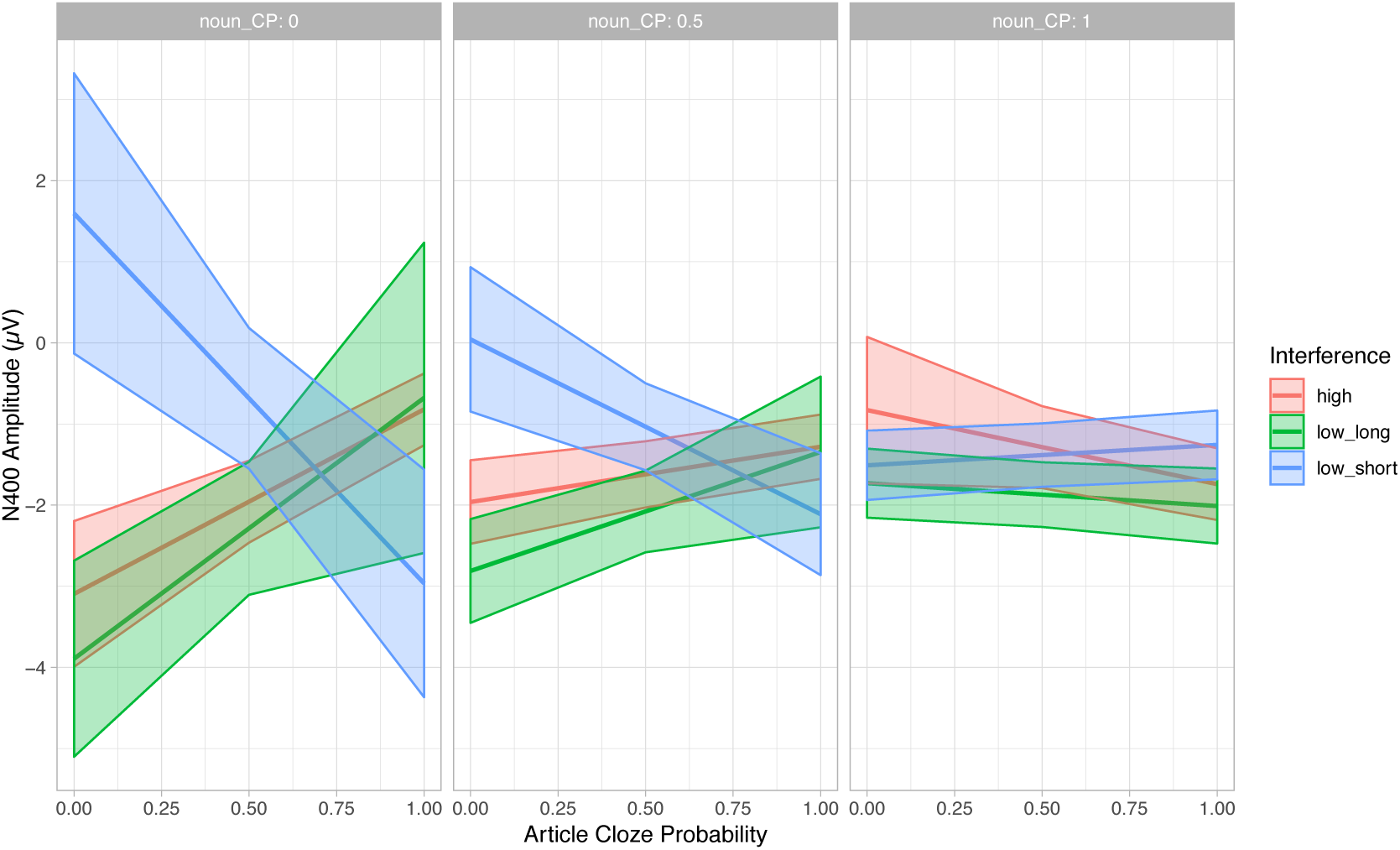
Predicted values of mean amplitude in the N400 time window at article position by interference, article CP and noun CP. Shaded regions indicate 95% confidence intervals.

**Table 2:** Article. N400 time window. Summary of parameters and random effects from the model produced by the call `lmer (formula = mean ~ 1 + sagittal * lateral * Interference * article_CP * noun_CP + scale(baseline) + (1 | subj) + (1 | item), data = det_n4, REML = F)`

| Linear mixed model fit by maximum likelihood [`lmerMod`] |  |  |  |  |
| --- | --- | --- | --- | --- |
| AIC | BIC | logLik | Deviance |  |
| 1392192 | 1393346 | -695984 | 1391968 | 220592 |
| Scaled residuals: |  |  |  |  |
| Min | 1Q | Median | 3Q | Max |
| -7.31 | -0.61 | 0.01 | 0.61 | 6.71 |
| Random effects: |  |  |  |  |
| Groups | Name | Variance | Std. Dev. |  |
| Item | (Intercept) | 0.3794 | 0.6159 |  |
| Subj | (Intercept) | 1.0862 | 1.0422 |  |
| Residual |  | 32.0418 | 5.6605 |  |
| Number of obs: 220704, groups: item, 78; subj, 32 |  |  |  |  |

**Table 3:** Article. N400 time window. Summary of fixed effects from the model produced by the call `lmer(formula = mean ~ 1 + sagittal * lateral * Interference * article_CP * noun_CP + scale(baseline) + (1 | subj) + (1 | item), data = det_n4, REML = F)`

|  | Estimate | Std. Error | t value |
| --- | --- | --- | --- |
| (Intercept) | -2.3 | 0.43 | -5.3 |
| sag[ant] | -1.2 | 0.45 | -2.6 |
| sag[post] | 1.1 | 0.47 | 2.5 |
| lat[left] | -1.1 | 0.45 | -2.5 |
| lat[right] | 1.2 | 0.45 | 2.6 |
| Interf[high] | -1.3 | 0.47 | -2.8 |
| Interf[low_long] | -2.1 | 0.46 | -4.5 |
| article_CP | 0.31 | 0.7 | 0.44 |
| noun_CP | 0.44 | 0.45 | 0.97 |
| scale(baseline) | 2.4 | 0.013 | 1.9e+02 |
| sag[ant]:lat[left] | 0.1 | 0.64 | 0.16 |
| sag[post]:lat[left] | -0.027 | 0.65 | -0.041 |
| sag[ant]:lat[right] | -0.35 | 0.64 | -0.55 |
| sag[post]:lat[right] | 0.12 | 0.65 | 0.19 |
| sag[ant]:Interf[high] | -0.096 | 0.53 | -0.18 |
| sag[post]:Interf[high] | 0.0058 | 0.55 | 0.01 |
| sag[ant]:Interf[low_long] | -0.11 | 0.61 | -0.19 |
| sag[post]:Interf[low_long] | 0.24 | 0.63 | 0.38 |
| lat[left]:Interf[high] | 0.58 | 0.53 | 1.1 |
| lat[right]:Interf[high] | -0.032 | 0.53 | -0.061 |
| lat[left]:Interf[low_long] | 0.95 | 0.61 | 1.6 |
| lat[right]:Interf[low_long] | -1.2 | 0.61 | -1.9 |
| sag[ant]:article_CP | -1.4 | 0.8 | -1.7 |

Table 3: Continued: Article. N400 time window. Summary of fixed effects from the model produced by the call `lmer(formula = mean ~ 1 + sagittal * lateral * Interference * article_CP * noun_CP + scale(baseline) + (1 | subj) + (1 | item), data = det_n4, REML = F)`
|  | Estimate | Std. Error | t value |
| --- | --- | --- | --- |
| sag[post]:article_CP | 1 | 0.83 | 1.3 |
| lat[left]:article_CP | 1.1 | 0.8 | 1.3 |
| lat[right]:article_CP | -0.64 | 0.8 | -0.79 |
| Interf[high]:article_CP | 2 | 0.77 | 2.6 |
| Interf[low_long]:article_CP | 2.9 | 0.99 | 2.9 |
| sag[ant]:noun_CP | 0.88 | 0.54 | 1.6 |
| sag[post]:noun_CP | -1.1 | 0.56 | -1.9 |
| lat[left]:noun_CP | 1.2 | 0.54 | 2.3 |
| lat[right]:noun_CP | -1.1 | 0.54 | -2 |
| Interf[high]:noun_CP | 1.8 | 0.66 | 2.8 |
| Interf[low_long]:noun_CP | 1.7 | 0.54 | 3.2 |
| article_CP:noun_CP | -0.62 | 0.8 | -0.78 |
| sag[ant]:lat[left]:Interf[high] | 0.29 | 0.76 | 0.38 |
| sag[post]:lat[left]:Interf[high] | -0.21 | 0.77 | -0.28 |
| sag[ant]:lat[right]:Interf[high] | 0.28 | 0.76 | 0.37 |
| sag[post]:lat[right]:Interf[high] | 0.028 | 0.77 | 0.036 |
| sag[ant]:lat[left]:Interf[low_long] | -0.41 | 0.87 | -0.48 |
| sag[post]:lat[left]:Interf[low_long] | 0.034 | 0.88 | 0.039 |
| sag[ant]:lat[right]:Interf[low_long] | 0.69 | 0.87 | 0.8 |
| sag[post]:lat[right]:Interf[low_long] | -0.62 | 0.88 | -0.7 |
| sag[ant]:lat[left]:article_CP | -0.67 | 1.1 | -0.58 |
| sag[post]:lat[left]:article_CP | 0.46 | 1.2 | 0.4 |
| sag[ant]:lat[right]:article_CP | 0.4 | 1.1 | 0.35 |
| sag[post]:lat[right]:article_CP | -0.041 | 1.2 | -0.035 |
| sag[ant]:Interf[high]:article_CP | 1.7 | 0.88 | 1.9 |
| sag[post]:Interf[high]:article_CP | -1.5 | 0.91 | -1.6 |
| sag[ant]:Interf[low_long]:article_CP | -0.66 | 1.3 | -0.52 |
| sag[post]:Interf[low_long]:article_CP | 0.46 | 1.3 | 0.35 |
| lat[left]:Interf[high]:article_CP | -0.66 | 0.88 | -0.75 |
| lat[right]:Interf[high]:article_CP | -0.13 | 0.88 | -0.14 |
| lat[left]:Interf[low_long]:article_CP | -0.56 | 1.3 | -0.44 |
| lat[right]:Interf[low_long]:article_CP | 2 | 1.3 | 1.6 |
| sag[ant]:lat[left]:noun_CP | -1 | 0.77 | -1.3 |
| sag[post]:lat[left]:noun_CP | 0.64 | 0.78 | 0.82 |
| sag[ant]:lat[right]:noun_CP | 0.6 | 0.77 | 0.78 |
| sag[post]:lat[right]:noun_CP | -0.27 | 0.78 | -0.34 |
| sag[ant]:Interf[high]:noun_CP | 0.97 | 0.75 | 1.3 |
| sag[post]:Interf[high]:noun_CP | -0.75 | 0.77 | -0.97 |
| sag[ant]:Interf[low_long]:noun_CP | -0.35 | 0.71 | -0.49 |
| sag[post]:Interf[low_long]:noun_CP | 0.15 | 0.73 | 0.21 |
| lat[left]:Interf[high]:noun_CP | -0.13 | 0.75 | -0.17 |
| lat[right]:Interf[high]:noun_CP | -0.79 | 0.75 | -1.1 |
| lat[left]:Interf[low_long]:noun_CP | -1.1 | 0.71 | -1.6 |
| lat[right]:Interf[low_long]:noun_CP | 1.7 | 0.71 | 2.4 |

Table 3: Continued: Article. N400 time window. Summary of fixed effects from the model produced by the call `lmer(formula = mean ~ 1 + sagittal * lateral * Interference * article_CP * noun_CP + scale(baseline) + (1 | subj) + (1 | item), data = det_n4, REML = F)`
|  | Estimate | Std. Error | t value |
| --- | --- | --- | --- |
| sag[ant]:article_CP:noun_CP | 0.85 | 0.92 | 0.93 |
| sag[post]:article_CP:noun_CP | -0.61 | 0.95 | -0.64 |
| lat[left]:article_CP:noun_CP | -1.3 | 0.92 | -1.4 |
| lat[right]:article_CP:noun_CP | 1.1 | 0.92 | 1.2 |
| Interf[high]:article_CP:noun_CP | -2.6 | 0.97 | -2.7 |
| Interf[low_long]:article_CP:noun_CP | -2.9 | 1.1 | -2.6 |
| sag[ant]:lat[left]:Interf[high]:article_CP | -0.33 | 1.3 | -0.26 |
| sag[post]:lat[left]:Interf[high]:article_CP | 0.23 | 1.3 | 0.18 |
| sag[ant]:lat[right]:Interf[high]:article_CP | -0.17 | 1.3 | -0.14 |
| sag[post]:lat[right]:Interf[high]:article_CP | -0.35 | 1.3 | -0.28 |
| sag[ant]:lat[left]:Interf[low_long]:article_CP | 0.83 | 1.8 | 0.46 |
| sag[post]:lat[left]:Interf[low_long]:article_CP | -0.35 | 1.8 | -0.19 |
| sag[ant]:lat[right]:Interf[low_long]:article_CP | -1.4 | 1.8 | -0.77 |
| sag[post]:lat[right]:Interf[low_long]:article_CP | 1.2 | 1.8 | 0.64 |
| sag[ant]:lat[left]:Interf[high]:noun_CP | -0.91 | 1.1 | -0.86 |
| sag[post]:lat[left]:Interf[high]:noun_CP | 0.5 | 1.1 | 0.47 |
| sag[ant]:lat[right]:Interf[high]:noun_CP | 0.056 | 1.1 | 0.053 |
| sag[post]:lat[right]:Interf[high]:noun_CP | -0.067 | 1.1 | -0.062 |
| sag[ant]:lat[left]:Interf[low_long]:noun_CP | 0.89 | 1 | 0.89 |
| sag[post]:lat[left]:Interf[low_long]:noun_CP | -0.27 | 1 | -0.26 |
| sag[ant]:lat[right]:Interf[low_long]:noun_CP | -0.99 | 1 | -0.98 |
| sag[post]:lat[right]:Interf[low_long]:noun_CP | 0.68 | 1 | 0.67 |
| sag[ant]:lat[left]:article_CP:noun_CP | 1.3 | 1.3 | 0.97 |
| sag[post]:lat[left]:article_CP:noun_CP | -0.87 | 1.3 | -0.65 |
| sag[ant]:lat[right]:article_CP:noun_CP | -0.62 | 1.3 | -0.47 |
| sag[post]:lat[right]:article_CP:noun_CP | 0.0008 | 1.3 | 0.0006 |
| sag[ant]:Interf[high]:article_CP:noun_CP | -2.7 | 1.1 | -2.4 |
| sag[post]:Interf[high]:article_CP:noun_CP | 2.4 | 1.2 | 2.1 |
| sag[ant]:Interf[low_long]:article_CP:noun_CP | 1.2 | 1.4 | 0.81 |
| sag[post]:Interf[low_long]:article_CP:noun_CP | -0.95 | 1.5 | -0.64 |
| lat[left]:Interf[high]:article_CP:noun_CP | -0.08 | 1.1 | -0.072 |
| lat[right]:Interf[high]:article_CP:noun_CP | 1.2 | 1.1 | 1.1 |
| lat[left]:Interf[low_long]:article_CP:noun_CP | 1 | 1.4 | 0.73 |
| lat[right]:Interf[low_long]:article_CP:noun_CP | -3 | 1.4 | -2.1 |
| sag[ant]:lat[left]:Interf[high]:article_CP:noun_CP | 0.96 | 1.6 | 0.6 |
| sag[post]:lat[left]:Interf[high]:article_CP:noun_CP | -0.43 | 1.6 | -0.27 |
| sag[ant]:lat[right]:Interf[high]:article_CP:noun_CP | -0.27 | 1.6 | -0.17 |
| sag[post]:lat[right]:Interf[high]:article_CP:noun_CP | 0.41 | 1.6 | 0.26 |
| sag[ant]:lat[left]:Interf[low_long]:article_CP:noun_CP | -1.6 | 2 | -0.77 |
| sag[post]:lat[left]:Interf[low_long]:article_CP:noun_CP | 0.68 | 2.1 | 0.33 |
| sag[ant]:lat[right]:Interf[low_long]:article_CP:noun_CP | 1.9 | 2 | 0.92 |
| sag[post]:lat[right]:Interf[low_long]:article_CP:noun_CP | -1.3 | 2.1 | -0.63 |

**Table 4:**
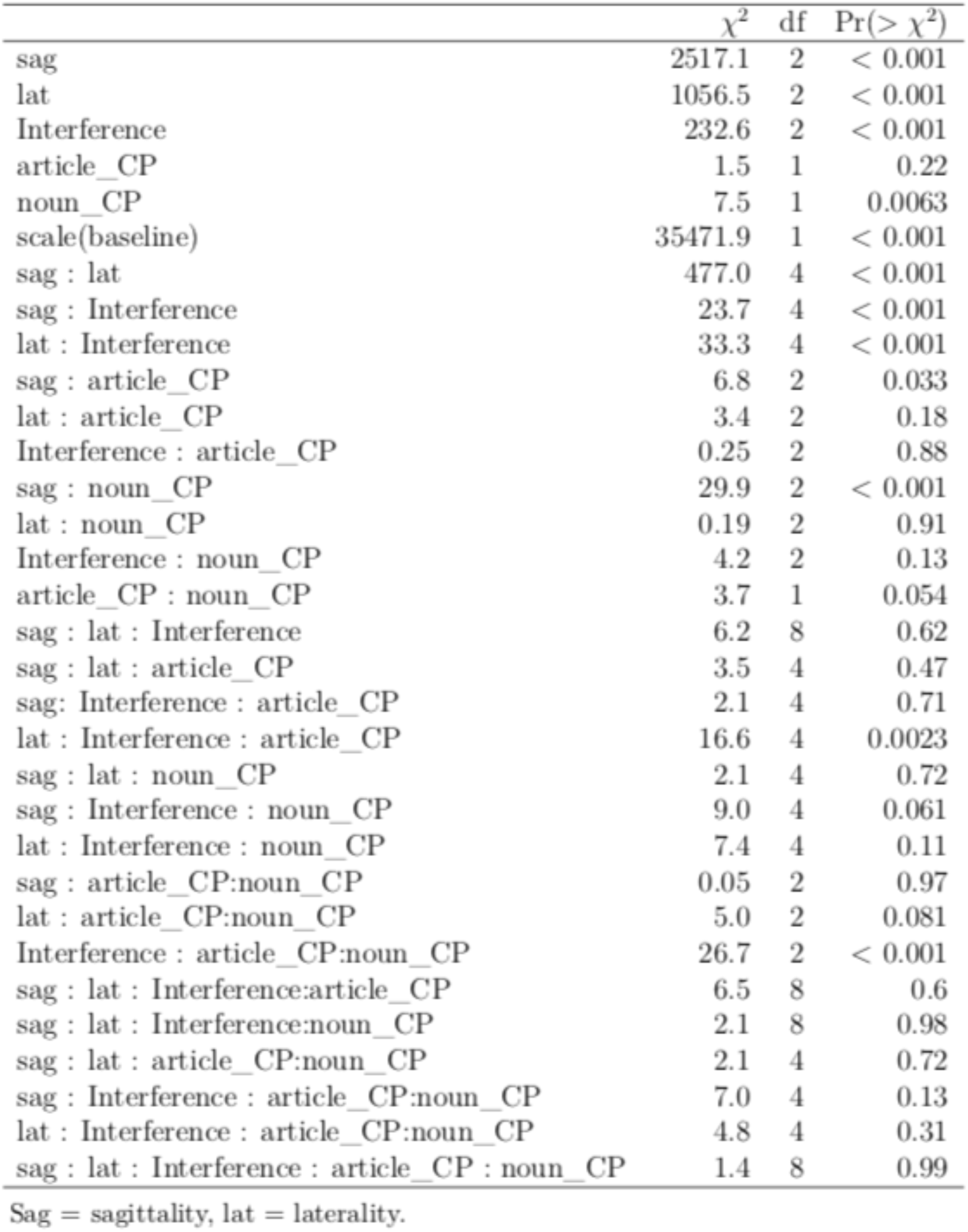
Summary of effects in the N400 time window after article onset (Type II Wald Test).

Effects of interference emerged only when both Article CP and Noun CP were low and manifested themselves in the form of increased negativity for the high interference and low interference / long target distance conditions in comparison to the low interference / short target distance condition. Table 5 shows the estimated marginal means of this interaction.

**Table 5:** Estimated marginal means for the Article in the N4(X) time window produced by the call `emmeans(det_n4_Ionly, c(“Interference”, “article_CP”, “noun_CP”), at = list(article_CP = c(0, 0.5, l),noun_CP = c(0, 0.5, 1)))`

| Interference | article_CP | noun_CP | emmean | SE | df | asympt.LCL | asympt.UCL |
| --- | --- | --- | --- | --- | --- | --- | --- |
| high | 0.0 | 0.0 | -3.581 | 0.459 | Inf | -4.479 | -2.682 |
| low_long | 0.0 | 0.0 | -4.379 | 0.617 | Inf | -5.588 | -3.171 |
| low_short | 0.0 | 0.0 | 1.110 | 0.881 | Inf | -0.616 | 2.837 |
| high | 0.5 | 0.0 | -2.443 | 0.258 | Inf | -2.948 | -1.937 |
| low_long | 0.5 | 0.0 | -2.771 | 0.418 | Inf | -3.591 | -1.951 |
| low_short | 0.5 | 0.0 | -1.170 | 0.443 | Inf | -2.037 | -0.303 |
| high | 1.0 | 0.0 | -1.305 | 0.226 | Inf | -1.747 | -0.863 |
| low_long | 1.0 | 0.0 | -1.163 | 0.975 | Inf | -3.075 | 0.749 |
| low_short | 1.0 | 0.0 | -3.451 | 0.716 | Inf | -4.854 | -2.047 |
| high | 0.0 | 0.5 | -2.447 | 0.263 | Inf | -2.962 | -1.933 |
| low_long | 0.0 | 0.5 | -3.297 | 0.326 | Inf | -3.937 | -2.658 |
| low_short | 0.0 | 0.5 | -0.442 | 0.454 | Inf | -1.332 | 0.448 |
| high | 0.5 | 0.5 | -2.107 | 0.208 | Inf | -2.515 | -1.698 |
| low_long | 0.5 | 0.5 | -2.564 | 0.257 | Inf | -3.067 | -2.060 |
| low_short | 0.5 | 0.5 | -1.520 | 0.274 | Inf | -2.056 | -0.984 |
| high | 1.0 | 0.5 | -1.766 | 0.202 | Inf | -2.162 | -1.369 |
| low_long | 1.0 | 0.5 | -1.830 | 0.474 | Inf | -2.759 | -0.901 |
| low_short | 1.0 | 0.5 | -2.597 | 0.384 | Inf | -3.349 | -1.845 |
| high | 0.0 | 1.0 | -1.314 | 0.460 | Inf | -2.216 | -0.412 |
| low_long | 0.0 | 1.0 | -2.215 | 0.217 | Inf | -2.641 | -1.790 |
| low_short | 0.0 | 1.0 | -1.995 | 0.218 | Inf | -2.422 | -1.568 |
| high | 0.5 | 1.0 | -1.771 | 0.258 | Inf | -2.275 | -1.266 |
| low_long | 0.5 | 1.0 | -2.356 | 0.204 | Inf | -2.755 | -1.957 |
| low_short | 0.5 | 1.0 | -1.869 | 0.199 | Inf | -2.260 | -1.478 |
| high | 1.0 | 1.0 | -2.227 | 0.225 | Inf | -2.669 | -1.786 |
| low_long | 1.0 | 1.0 | -2.497 | 0.236 | Inf | -2.959 | -2.034 |
| low_short | 1.0 | 1.0 | -1.743 | 0.216 | Inf | -2.167 | -1.319 |

Results are averaged over the levels of: sagittal, lateral
Degrees-of-freedom method: asymptotic
Confidence level used: 0.95.

It was mentioned in the Methods section, that the combinations of CP values are not evenly distributed across conditions; therefore we present the data basis for the above-described results in Figure 4. Across the Noun CP scale, for the high interference conditions, the Article CP values cover the range from approx. 0.5 to 1. Additionally, for the low interference conditions, Noun CP ranged from approx. 0.5 to 1. See appendix, for the results at the positon of the noun.

**Figure 4:**
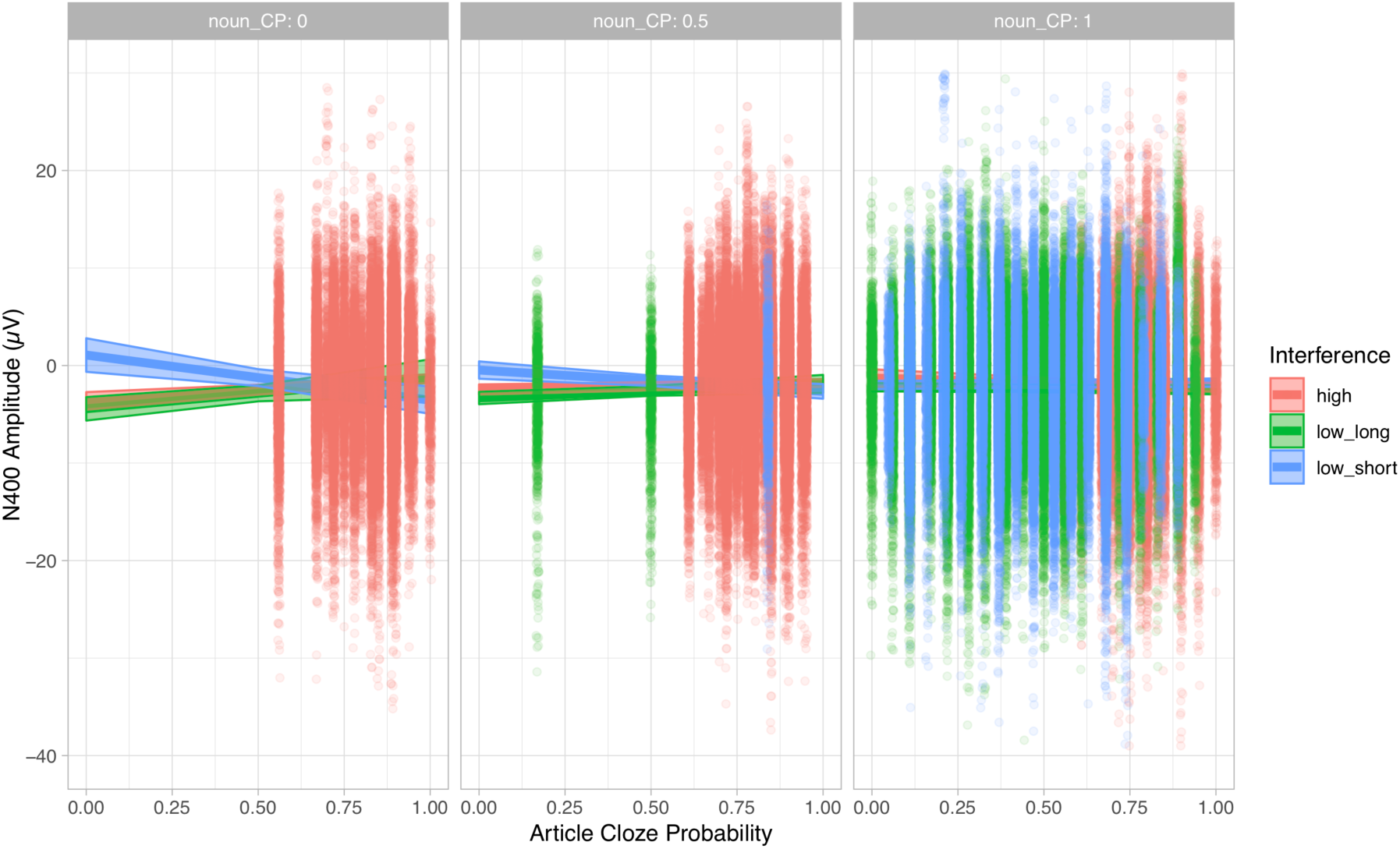
Distribution of data points with estimated effects at article position in the background.

## 4. Discussion

We have presented an ERP study that investigated the interplay of prediction and interference within sentence pairs. In addition to the experimental manipulations, we included offline cloze probability (CP) values of the critical article and following noun in linear mixed model analyses of mean single trial EEG activity within the N400 time window (300 – 500 ms post stimulus onset). At the position of the critical article, the linear mixed model estimated a negativity for conditions with a high-interference distractor and for conditions with a close, low-interference distractor (i.e. long distance to target) in comparison to conditions with a distant, low-interference distractor (i.e. short distance to target). These effects only emerged when the CP of the article itself and of the following noun were low.

Before we discuss the results, we recapitulate what the different combinations of CP values mean for the processing of our critical article (see Figure 6). The combination of a low cloze article and a low cloze noun represents cases where the article was not predicted based on the context and even after article processing the noun did not become predictable. This combination was not observed in our materials (see Figure 2), hence results for that combination are solely based on model estimates. In contrast, the combination of a high cloze article and a high cloze noun represents cases where the whole target NP was predicted before article presentation and was already confirmed by the matching article. This combination is seen for all our conditions. The combination of a low cloze article and a high cloze noun represents that the critical NP was not predicted before the article was processed but its processing led to a clear noun prediction, hence the noun prediction is quite recent. This combination is seen only for low interference conditions. The combination of a high cloze article and a low cloze noun represents that the article was predicted but after its processing the target noun was not predicted. Intuitively, this can only be the case in high interference conditions, when the distractor is predicted to occur in the target sentence. Then, while the distractor article was predicted, because it shares the gender feature with the target article, its CP is high, but the CP of the target noun is low. The combination of a high cloze article and a low cloze noun for the low interference conditions did not occur in our materials (see Figure 2), therefore the results for this combination in the low interference conditions are based on the model estimates only.

**Figure 6:**
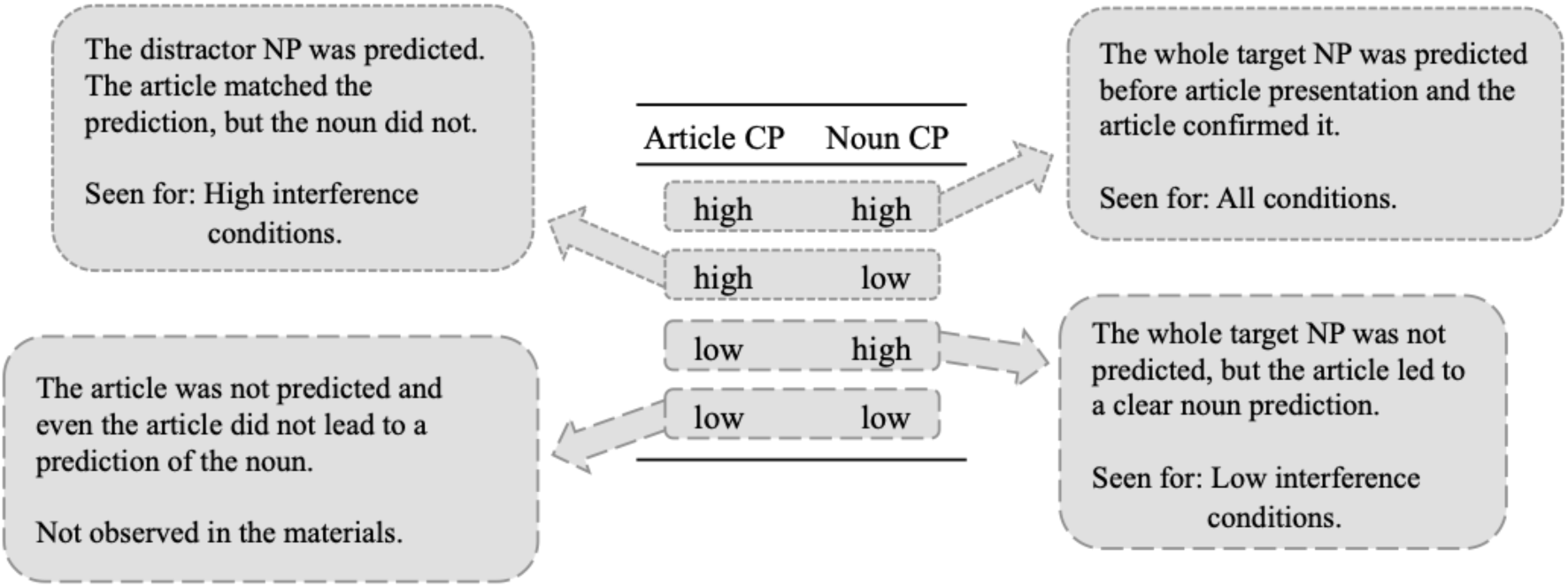
Visualization what the different combinations of CP values mean for the processing of our critical article and the following noun.

In the following, we will first discuss the results and subsequently, we will consider potential theoretical implications of our findings, before offering some conclusions.

### 4.1 Article

Effects at the article emerged only when the CP of both the article and noun were low. Thus, under conditions of high predictability, the distractors were essentially “ignored” by the processing system. We suggest that this aligns well with a predictive coding perspective, according to which high predictability leads to all (or almost all) of the sensory input for a current item being “explained away” (cf. Friston, 2010; and see the introduction for further discussion).

In contrast, when predictability is low, bottom-up information gains more weight. Since the input is not “explained away” under these circumstances, bottom-up sensory information (corresponding to prediction error), is propagated up through the levels of the hierarchically organized predictive processing architecture. Prediction errors induce model updating, which – we assume – requires an evaluation of all (other) possible candidate referents for the article and upcoming noun. Since this evaluation necessarily involves retrieval of the candidates, it is a natural locus for interference. We propose here that interference modulates the complexity of the model update and, thereby, N400 amplitude. In sentences of the type employed in our experiment, the required model update involves retrieving the correct referent for the upcoming noun. This retrieval operation is facilitated under conditions of low interference: since the gender information provided by the article uniquely matches exactly one noun in the context, this scenario involves a low risk of prediction errors according to the next word, i.e. the noun, in spite of the fact that the CP of noun and article was low prior to processing of the corresponding stimuli. In essence, retrieval is easy due to the lack of alternatives – possibly due to the restricted experimental context. In the low interference / short target distance condition, retrieval of the target noun is additionally supported by recency (see section 2.3 for a discussion of recency effects in our materials as revealed by the pretest). The target noun was introduced recently and is neuronally represented via short-term synaptic plasticity. The gender retrieval cue uniquely matches with parts of that recently active neuronal assembly and re-instantiates its firing pattern without conflict (therefore, we found no effect of proactive interference in these conditions).

In the low interference / long target distance condition, the target noun was processed some time ago and is again represented via synaptic weights. But in contrast to the low interference / short target distant condition, in this condition the distractor was processed after the target. The distractor’s firing pattern probably includes neurons that represent the target, therefore disturbing the synaptic weights that carry the target representation (retroactive interference). Thus, during retrieval, the target representation is weaker than the recently established distractor representation and they additionally overlap to some degree, which may result in retrieval of both nouns (despite only one of them matching the gender retrieval cue). This renders model updating more complex, due to the two conflicting information sources, and we observe a more pronounced N400 effect under these conditions.

Finally, under conditions of high interference, a low CP article induces a prediction error but, at the same time, provides no additional evidence to guide a model update. The gender retrieval cue matches with two overlapping neuronal assemblies that represent the two nouns that have been introduced in the context sentence. Presumably, both firing patterns are reactivated in this case. While a model update is clearly required here, it is associated with a high degree of uncertainty, again correlating with a more pronounced N400 amplitude.

Thus, in line with Bornkessel-Schlesewsky and Schlesewsky’s (2019) proposal, our results at the position of the critical article suggest that N400 amplitude during incremental sentence comprehension does not reflect the presence / absence of a prediction error per se, but rather the effects of a prediction error on internal model updating. The present findings suggest that interference may play an important role in this regard, as the presence of interfering distractors not only impacts memory retrieval (i.e. the likelihood of retrieving the correct item from memory), but also the complexity of model updating. In addition, if an account along these lines is correct, it suggests that, prediction error within a hierarchically organized predictive coding architecture might trigger memory retrieval. We return to this point in section 4.2 below.

### 4.2 Theoretical implications

Our work has potential implications for all of the theoretical frameworks under discussion: neuronal memory representation, predictive coding, cue-based parsing (retrieval) and the functional interpretation of N400 amplitude as reflecting model updating via precision-weighted prediction error signals. We will briefly discuss each of these in turn and finally, point out our contribution to the debate of form-based predictions during language comprehension.

According to Jonides et al.’s (2008) proposal regarding the neuronal mechanisms of memory, an item within the attentional focus is represented with an active firing pattern. When the item is no longer in focus, its representation is carried by short-term synaptic plasticity. Perception, attention shifts, retrieval and prediction can all serve to reinstantiate the active firing pattern of the item, thus providing a neurobiologically grounded theoretical basis for the interplay of prediction and retrieval.

Within a hierarchically organized predictive coding architecture, prediction errors trigger model updating. Our results provide some initial indications as to how this updating process may be shaped by different information sources in the context of a complex domain such as higher-order language processing. Specifically, the updates required in our experimental conditions cannot depend entirely on incoming sensory information, but also require recourse to previous input, which needs to be accessed via retrieval operations. In neural terms, retrieval can be conceived as the reinstatement of an item’s firing pattern and we propose here that prediction errors play an important role in triggering this reinstatement. Specifically, the information carried by the error signal corresponds to the notion of a retrieval cue in cognitive terms. Retrieval can, in turn, be influenced by interference – overlap in the neural firing patterns associated with target and distractor items – which lowers the probability of correctly retrieving the target by reinstating its firing pattern or, in a worst-case scenario, may even favor retrieval of an incorrect (distractor) item. If the incorrect item is retrieved, the priors of the new model are incorrect, thereby inevitably leading to prediction error as the sentence continues. If interference manifests as multiple possible targets for the retrieval, new predictions that are based on this retrieval situation are less trusted because of the increased risk of prediction error. The resulting model thus has less reliability due to the lower-precision predictions. In summary, our findings suggest that: (a) prediction errors trigger memory retrieval when the incoming sensory information is not sufficient to guide model updating; and (b) retrieval interference during model updating diminishes the ability of the model update to minimize future prediction errors.

Under the cue-based parsing framework, memory representations not only comprise already processed words, but also predictions for upcoming words or constituents. The former are retrieved when new words need to be integrated with them (e.g. to establish co-reference or to saturate dependencies between heads such as verbs and their dependent arguments). By contrast, the latter are retrieved when the predicted words are actually encountered. The retrieval operation investigated with the presented study might be comparable to those during reflexive-/reciprocal-antecedent or ellipsis processing. Rather than retrieving an argument-verb dependency, for example, it calls for the retrieval of the correct (co-)referent for the current item. But there are no structural constraints (e.g. c-command in Chomskyan frameworks), which function as additional retrieval cues during the retrieval of our referent.

There appears to be a conceptual discrepancy between the memory representation of a predicted word that needs to be retrieved in cue-based parsing and predictions in the sense of predictive coding. In cue-based parsing, the memory representation of the representation is shifted out of attentional focus when other words are processed (see McElree, 2006). Thus, when the predicted word is encountered, the prediction must be retrieved in order to allow for an integration of the predicted, newly encountered word with the corresponding “old” word. Within the predictive coding architecture, by contrast, the prediction of an upcoming stimulus of a specific kind would be (repeatedly?) communicated via feedback connections until the stimulus is encountered, at which point the prediction would “explain away” the sensory input. However, given the propensity for non-adjacent dependencies in language, all stimuli that are encountered between the generation of the prediction and the predicted stimulus’ appearance would lead to prediction error. This could either engender model updating and subsequent dismissal of the prediction, or – what is perhaps more likely – to a generalization of the prediction (e.g. the prediction that a noun phrase should be followed, at some point, by a verb). This may lead to maintenance or even strengthening of the prediction (cf. anti-locality effects, (Husain, Vasishth, & Srinivasan, 2014; Konieczny, 2000)), which appears incompatible with the notion that a predicted memory representation disappears from the focus of attention and becomes active again later. One way to operationalize predictions for non-adjacent elements in a predictive coding framework may be to assume that these correspond to higher levels – and hence longer timescales – within the hierarchically organized predictive coding architecture (cf. Bornkessel-Schlesewsky & Schlesewsky, 2019; Bornkessel-Schlesewsky et al., 2015).

The N400-as-model-updating account by Bornkessel-Schlesewsky and Schlesewsky (2019) posits that N400 amplitude reflects precision-weighted prediction errors and, hence, the consequences of a prediction error for model updating rather than prediction-error-related activity per se. Initially, the notion of precision-weighting was introduced to account for N400 modulations via the relevance (cue validity) of a particular feature within a given language. The results of the present study provide evidence to suggest that this notion could be extended to varying validity of a cue according to the specific context. Interference contexts reduce the validity of a cue as already described by the term of cue diagnosticity (Nairne, 2002). A cue is diagnostic if it uniquely picks out the target and excludes the distractors (see Martin et al., 2012 for detailed discussion). Thus, interference during a retrieval operation as part of a model update is closely related to the precision of future predictions.

Finally, the present findings provide further converging support for the notion that predictive processing during language comprehension may – at least under certain circumstances – lead to the anticipation of features and thereby the *form* of upcoming input elements. This idea was prominently put forward by DeLong, Urbach, and Kutas (2005), who reported evidence for an anticipation of the English articles “a” versus “an” on the basis of a predicted, following noun. A recent large-scale replication study failed to replicate this effect at the position of the article (Nieuwland et al., 2018), thus leading to a controversial discussion of whether the language comprehension system indeed predicts upcoming form-based features. By showing effects of prediction and interference at an article (i.e. a function word that does not vary in its compatibility with the context unless viewed in conjunction with the following noun), our study supports the notion that feature- and thereby form-based predictions can indeed be generated. This adds further support to a growing body of literature in languages other than English, e.g. (noun-)gender-based N400 effects at the position of pre-nominal articles in spoken Spanish (Wicha, Bates, Moreno, & Kutas, 2003; but see Wicha, Moreno, & Kutas, 2004 for a P600 effect in a similar design) and pre-nominal adjectives in Polish (Szewczyk & Schriefers, 2013), as well as handshape-based N400 effects prior to the onset of a critical sign in German sign language (Hosemann, Herrmann, Steinbach, Bornkessel-Schlesewsky, & Schlesewsky, 2013). By further demonstrating that these pre-nominal prediction effects are modulated by a range of influences that have not been examined to date – most notably interference, but also the interaction of article CP and noun CP – our study provides an initial indication as to why previous studies may have shown apparently contradictory results.

## 5. Conclusion

The results of the present ERP study suggest that prediction outweighs interference. Under high predictability, interference effects do not arise, because the sensory input at the retrieval site is “explained away” in the sense of predictive coding. Hence, retrieval is not necessary, thus eliminating the possibility of interference. We conclude that interference may play a crucial role in a hierarchically organized, cortical predictive coding architecture for language as it influences the complexity of model updating and the precision of future predictions — particularly when predictability is low.

## Acknowledgements

The authors would like to thank Jonas Diekmann and Phillip M. Alday for helpful discussions related to the research reported here. We also thank Mante S. Nieuwland and two anonymous reviewers for their helpful comments on an earlier version of this manuscript. Thanks are also due to Anja Bergmair, Nadine Furtner and Veronica Baldin for their contributions to the experimental materials and to Barbara Sophie Hartl and again Anja Bergmair for their assistance during data acquistion. This research was supported by a research grant awarded to P.S. by the University of Salzburg. I.B.-S. acknowledges the support of an Australian Research Council Future Fellowship (FT160100437).

## Appendix

### A.1. Statistical Analysis

The noun was analyzed using linear mixed model analysis. It was carried out analog to the article analysis with the following expections: The two models differed in the coding of the factors interference and distance. In the noun model, we modeled the two factors separately: Interference (high, low) and Distance between target and retrieval site (short, long). The z-transformed baseline activity from 200 ms pre-stimulus until stimulus onset used in the noun analysis, corresponds to the N400 interval of the critical article.

**Figure S1:**
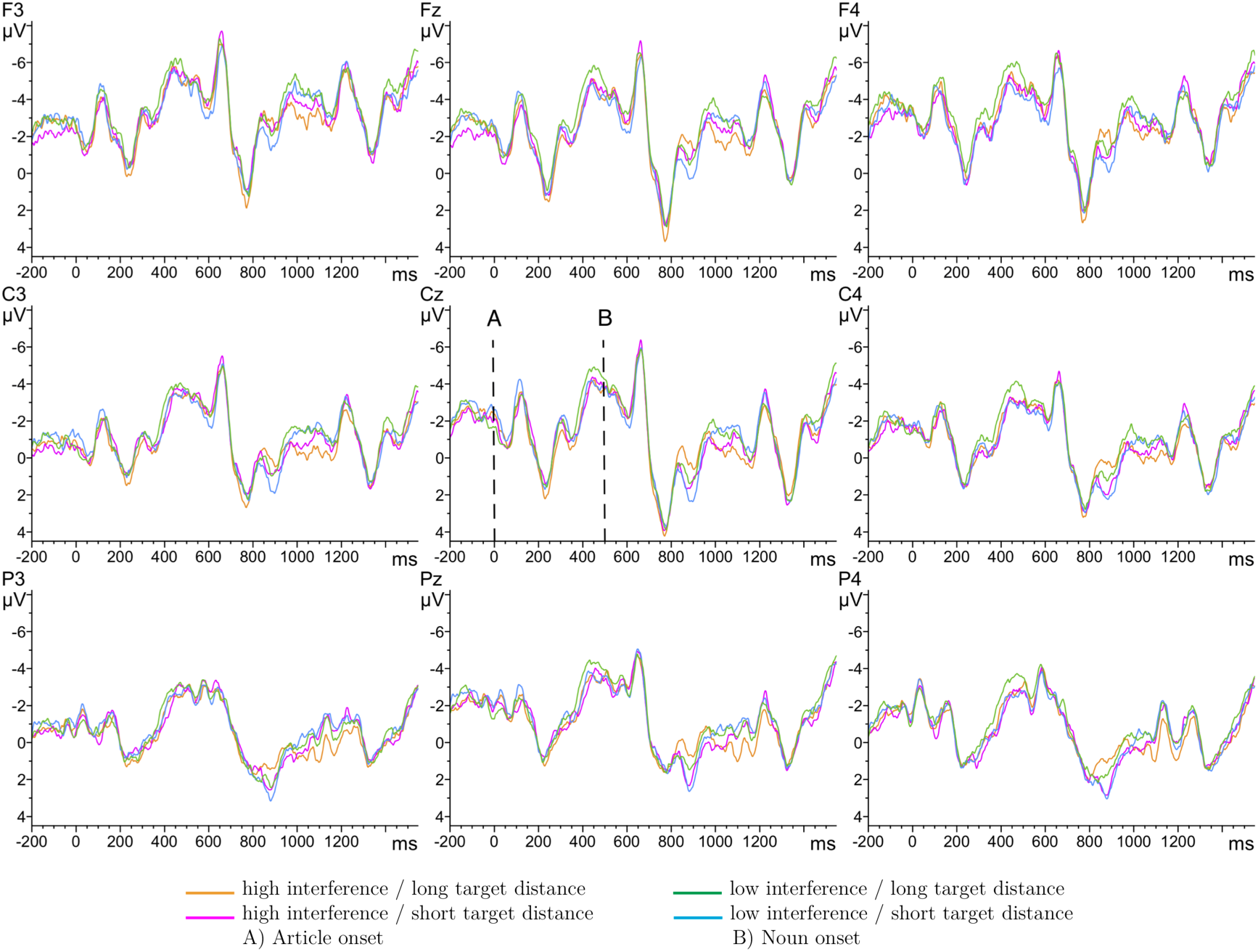
Grand averages at nine selected electrode sites for the critical article and following noun. A) Article onset, B) Noun onset. Negativity is plotted upwards.

### A.2. Electrophysiological Results

We present grand averages for the noun in Figure S1. The full summary for the linear mixed model for the noun is shown in Tables S1 and S2. Type-II Wald tests are shown in Table S3; we focus on the Interference x Target Distance x Article CP x Noun CP interaction (χ^2^(1) = 36.27, p < 0.001), which is visualized in Figure S1; the corresponding estimated marginal means are shown in Table S4. The absence of an interaction with sagittality indicates that the effects are broadly distributed. We do not further investigate the interaction of Laterality x Interference x Target Distance x Article CP x Noun CP (χ^2^(2) = 8.07, p = 0.018) because the meaning of laterality effects between ERPs in language comprehension is not clear. The effects pattern of the four-way interaction is complex and therefore to maintain intelligibility, we focus on the bigger differences between the confidence intervals.

**Table S1:** Summary of the random intercepts only model in the time window 300 - 500 ms from Noiui onset.

| Linear mixed model fit by maximum likelihood |  |  |  |  |
| --- | --- | --- | --- | --- |
| AIC | BIC | logLik | Deviance | df.resid. |
| 1399485 | 1401010 | -699595 | 1399189 | 219500 |
| Scaled residuals: |  |  |  |  |
| Min | 1Q | Median | 3Q | Max |
| -8.256 | -0.601 | 0.004 | 0.609 | 5.866 |
| Random effects: |  |  |  |  |
| Groups | Name | Variance | Std. Dev. |  |
| Item | (Intercept) | 0.682 | 0.826 |  |
| Subj | (Intercept) | 2.257 | 1.502 |  |
| Residual |  | 34.122 | 5.841 |  |
| Number of obs: 219648, groups: Item, 78; Subj, 32 |  |  |  |  |

**Table S2:** Noun. N400 time window. Summary of fixed effects from the model produced by the call `lmer(formula = mean ~ 1 + sagittal * lateral * Interference * Target_Distance * article_CP * noun_CP + scale(baseline) + (1 | subj) + (1 | item), data = noun_n4, REML = F)`. Sag = sagittal, lat = lateral, ant = anterior, post = posterior. Interf = Interference, TDist = Target_Distance, aCP = article_CP, nCP = noun_CP.

|  | Estimate | Std. Error | t value |
| --- | --- | --- | --- |
| (Intercept) | -3 | 0.48 | -6.1 |
| sag[ant] | -0.55 | 0.46 | -1.2 |
| sag[post] | 0.45 | 0.48 | 0.95 |
| lat[left] | 0.97 | 0.46 | 2.1 |
| lat[right] | -0.6 | 0.46 | -1.3 |
| Interf[high] | -0.96 | 0.4 | -2.4 |
| TDist[long] | 3.7 | 0.36 | 10 |
| aCP | 4.3 | 0.63 | 6.9 |
| nCP | 3.1 | 0.56 | 5.6 |
| scale(baseline) | 2.2 | 0.013 | 1.7e+02 |
| sag[ant]:lat[left] | -0.99 | 0.65 | -1.5 |
| sag[post]:lat[left] | 0.57 | 0.67 | 0.85 |
| sag[ant]:lat[right] | 0.1 | 0.65 | 0.16 |
| sag[post]:lat[right] | 0.073 | 0.67 | 0.11 |
| sag[ant]:Interf[high] | -0.25 | 0.46 | -0.54 |
| sag[post]:Interf[high] | 0.41 | 0.48 | 0.87 |
| lat[left]:Interf[high] | -0.4 | 0.46 | -0.86 |
| lat[right]:Interf[high] | 0.39 | 0.46 | 0.86 |
| sag[ant]:TDist[long] | 1.4 | 0.46 | 3.1 |
| sag[post]:TDist[long] | -1.1 | 0.48 | -2.4 |
| lat[left]:TDist[long] | -0.65 | 0.46 | -1.4 |
| lat[right]:TDist[long] | 0.48 | 0.46 | 1 |
| Interf[high]:TDist[long] | 0.95 | 0.37 | 2.6 |
| sag[ant]:aCP | 0.22 | 0.72 | 0.3 |
| sag[post]:aCP | -0.69 | 0.75 | -0.93 |

Table S2: Continued: Noun. N400 time window.
|  | Estimate | Std. Error | t value |
| --- | --- | --- | --- |
| lat[left]:aCP | -2.7 | 0.72 | -3.7 |
| lat[right]:aCP | 1.2 | 0.72 | 1.7 |
| Interf[high]:aCP | 0.06 | 0.63 | 0.095 |
| TDist[long]:aCP | -2.9 | 0.57 | -5.1 |
| sag[ant]:nCP | -0.67 | 0.67 | -1 |
| sag[post]:nCP | 0.14 | 0.69 | 0.21 |
| lat[left]:nCP | -1.4 | 0.67 | -2.2 |
| lat[right]:nCP | 0.76 | 0.67 | 1.1 |
| Interf[high]:nCP | 1.2 | 0.59 | 2.1 |
| TDist[long]:nCP | -5 | 0.52 | -9.6 |
| aCP:nCP | -5.2 | 0.83 | -6.3 |
| sag[ant]:lat[left]:Interf[high] | -1.2 | 0.65 | -1.8 |
| sag[post]:lat[left]:Interf[high] | 1.3 | 0.67 | 1.9 |
| sag[ant]:lat[right]:Interf[high] | 0.013 | 0.65 | 0.02 |
| sag[post]:lat[right]:Interf[high] | -0.43 | 0.67 | -0.64 |
| sag[ant]:lat[left]:TDist[long] | 0.61 | 0.65 | 0.94 |
| sag[post]:lat[left]:TDist[long] | -0.53 | 0.67 | -0.8 |
| sag[ant]:lat[right]:TDist[long] | -0.24 | 0.65 | -0.36 |
| sag[post]:lat[right]:TDist[long] | 0.17 | 0.67 | 0.25 |
| sag[ant]:Interf[high]:TDist[long] | -1.3 | 0.46 | -2.7 |
| sag[post]:Interf[high]:TDist[long] | 0.98 | 0.48 | 2.1 |
| lat[left]:Interf[high]:TDist[long] | -0.92 | 0.46 | -2 |
| lat[right]:Interf[high]:TDist[long] | 0.2 | 0.46 | 0.44 |
| sag[ant]:lat[left]:aCP | 0.098 | 1 | 0.095 |
| sag[post]:lat[left]:aCP | 0.17 | 1 | 0.16 |
| sag[ant]:lat[right]:aCP | 0.71 | 1 | 0.69 |
| sag[post]:lat[right]:aCP | -0.91 | 1 | -0.87 |
| sag[ant]:Interf[high]:aCP | -0.3 | 0.72 | -0.41 |
| sag[post]:Interf[high]:aCP | 0.31 | 0.75 | 0.42 |
| lat[left]:Interf[high]:aCP | 1.4 | 0.72 | 2 |
| lat[right]:Interf[high]:aCP | -0.85 | 0.72 | -1.2 |
| sag[ant]:TDist[long]:aCP | 0.016 | 0.72 | 0.022 |
| sag[post]:TDist[long]:aCP | -0.17 | 0.75 | -0.23 |
| lat[left]:TDist[long]:aCP | 0.86 | 0.72 | 1.2 |
| lat[right]:TDist[long]:aCP | -1.8 | 0.72 | -2.6 |
| Interf[high]:TDist[long]:aCP | -3 | 0.57 | -5.2 |
| sag[ant]:lat[left]:nCP | 0.86 | 0.95 | 0.91 |
| sag[post]:lat[left]:nCP | -0.58 | 0.96 | -0.6 |
| sag[ant]:lat[right]:nCP | 0.26 | 0.95 | 0.27 |
| sag[post]:lat[right]:nCP | -0.2 | 0.96 | -0.2 |
| sag[ant]:Interf[high]:nCP | 0.41 | 0.67 | 0.62 |
| sag[post]:Interf[high]:nCP | -0.8 | 0.69 | -1.2 |

Table S2: Continued: Noun. N400 time window.
|  | Estimate | Std. Error | t value |
| --- | --- | --- | --- |
| lat[left]:Interf[high]:nCP | 0.48 | 0.67 | 0.72 |
| lat[right]:Interf[high]:nCP | -0.39 | 0.67 | -0.59 |
| sag[ant]:TDist[long]:nCP | -1.5 | 0.67 | -2.3 |
| sag[post]:TDist[long]:nCP | 1.3 | 0.69 | 1.8 |
| lat[left]:TDist[long]:nCP | 1.4 | 0.67 | 2 |
| lat[right]:TDist[long]:nCP | -0.96 | 0.67 | -1.4 |
| Interf[high]:TDist[long]:nCP | -2 | 0.53 | -3.9 |
| sag[ant]:aCP:nCP | -0.57 | 0.96 | -0.6 |
| sag[post]:aCP:nCP | 1.3 | 0.99 | 1.3 |
| lat[left]:aCP:nCP | 2.8 | 0.96 | 2.9 |
| lat[right]:aCP:nCP | -1.1 | 0.96 | -1.2 |
| Interf[high]:aCP:nCP | -0.091 | 0.83 | -0.11 |
| TDist[long]:aCP:nCP | 4.2 | 0.75 | 5.6 |
| sag[ant]:lat[left]:Interf[high]:TDist[long] | -0.4 | 0.65 | -0.62 |
| sag[post]:lat[left]:Interf[high]:TDist[long] | 0.63 | 0.67 | 0.95 |
| sag[ant]:lat[right]:Interf[high]:TDist[long] | 1.5 | 0.65 | 2.2 |
| sag[post]:lat[right]:Interf[high]:TDist[long] | -1.2 | 0.67 | -1.7 |
| sag[ant]:lat[left]:Interf[high]:aCP | 2 | 1 | 1.9 |
| sag[post]:lat[left]:Interf[high]:aCP | -2 | 1 | -1.9 |
| sag[ant]:lat[right]:Interf[high]:aCP | -0.66 | 1 | -0.65 |
| sag[post]:lat[right]:Interf[high]:aCP | 1.1 | 1 | 1.1 |
| sag[ant]:lat[left]:TDist[long]:aCP | -0.93 | 1 | -0.9 |
| sag[post]:lat[left]:TDist[long]:aCP | 0.87 | 1 | 0.83 |
| sag[ant]:lat[right]:TDist[long]:aCP | 0.42 | 1 | 0.41 |
| sag[post]:lat[right]:TDist[long]:aCP | -0.059 | 1 | -0.056 |
| sag[ant]:Interf[high]:TDist[long]:aCP | -0.23 | 0.72 | -0.32 |
| sag[post]:Interf[high]:TDist[long]:aCP | 0.37 | 0.75 | 0.5 |
| lat[left]:Interf[high]:TDist[long]:aCP | 1.2 | 0.72 | 1.7 |
| lat[right]:Interf[high]:TDist[long]:aCP | 0.94 | 0.72 | 1.3 |
| sag[ant]:lat[left]:Interf[high]:nCP | 1.7 | 0.95 | 1.8 |
| sag[post]:lat[left]:Interf[high]:nCP | -1.7 | 0.96 | -1.8 |
| sag[ant]:lat[right]:Interf[high]:nCP | 0.071 | 0.95 | 0.075 |
| sag[post]:lat[right]:Interf[high]:nCP | 0.63 | 0.96 | 0.65 |
| sag[ant]:lat[left]:TDist[long]:nCP | -0.82 | 0.95 | -0.86 |
| sag[post]:lat[left]:TDist[long]:nCP | 0.62 | 0.96 | 0.64 |
| sag[ant]:lat[right]:TDist[long]:nCP | -0.52 | 0.95 | -0.54 |
| sag[post]:lat[right]:TDist[long]:nCP | 0.46 | 0.96 | 0.47 |
| sag[ant]:Interf[high]:TDist[long]:nCP | 1.5 | 0.67 | 2.3 |
| sag[post]:Interf[high]:TDist[long]:nCP | -1.1 | 0.69 | -1.6 |
| lat[left]:Interf[high]:TDist[long]:nCP | 1.8 | 0.67 | 2.6 |
| lat[right]:Interf[high]:TDist[long]:nCP | -0.8 | 0.67 | -1.2 |
| sag[ant]:lat[left]:aCP:nCP | -0.54 | 1.4 | -0.4 |

Table S2: Continued: Noun. N400 time window.
|  | Estimate | Std. Error | t value |
| --- | --- | --- | --- |
| sag[post]:lat[left]:aCP:nCP | 0.29 | 1.4 | 0.21 |
| sag[ant]:lat[right]:aCP:nCP | -0.85 | 1.4 | -0.63 |
| sag[post]:lat[right]:aCP:nCP | 0.71 | 1.4 | 0.51 |
| sag[ant]:Interf[high]:aCP:nCP | 0.23 | 0.96 | 0.24 |
| sag[post]:Interf[high]:aCP:nCP | 0.049 | 0.99 | 0.05 |
| lat[left]:Interf[high]:aCP:nCP | -1.5 | 0.96 | -1.6 |
| lat[right]:Interf[high]:aCP:nCP | 0.75 | 0.96 | 0.79 |
| sag[ant]:TDist[long]:aCP:nCP | -0.072 | 0.96 | -0.075 |
| sag[post]:TDist[long]:aCP:nCP | 0.25 | 0.99 | 0.26 |
| lat[left]:TDist[long]:aCP:nCP | -1.6 | 0.96 | -1.7 |
| lat[right]:TDist[long]:aCP:nCP | 2.5 | 0.96 | 2.6 |
| Interf[high]:TDist[long]:aCP:nCP | 4.6 | 0.75 | 6 |
| sag[ant]:lat[left]:Interf[high]:TDist[long]:aCP | 0.74 | 1 | 0.72 |
| sag[post]:lat[left]:Interf[high]:TDist[long]:aCP | -1.1 | 1 | -1 |
| sag[ant]:lat[right]:Interf[high]:TDist[long]:aCP | -2.1 | 1 | -2 |
| sag[post]:lat[right]:Interf[high]:TDist[long]:aCP | 1.4 | 1 | 1.4 |
| sag[ant]:lat[left]:Interf[high]:TDist[long]:nCP | 0.31 | 0.95 | 0.33 |
| sag[post]:lat[left]:Interf[high]:TDist[long]:nCP | -0.72 | 0.96 | -0.74 |
| sag[ant]:lat[right]:Interf[high]:TDist[long]:nCP | -2.4 | 0.95 | -2.5 |
| sag[post]:lat[right]:Interf[high]:TDist[long]:nCP | 1.9 | 0.96 | 2 |
| sag[ant]:lat[left]:Interf[high]:aCP:nCP | -2.6 | 1.4 | -1.9 |
| sag[post]:lat[left]:Interf[high]:aCP:nCP | 2.5 | 1.4 | 1.8 |
| sag[ant]:lat[right]:Interf[high]:aCP:nCP | 0.64 | 1.4 | 0.47 |
| sag[post]:lat[right]:Interf[high]:aCP:nCP | -1.4 | 1.4 | -1 |
| sag[ant]:lat[left]:TDist[long]:aCP:nCP | 1.1 | 1.4 | 0.84 |
| sag[post]:lat[left]:TDist[long]:aCP:nCP | -1 | 1.4 | -0.74 |
| sag[ant]:lat[right]:TDist[long]:aCP:nCP | 0.53 | 1.4 | 0.39 |
| sag[post]:lat[right]:TDist[long]:aCP:nCP | -0.75 | 1.4 | -0.54 |
| sag[ant]:Interf[high]:TDist[long]:aCP:nCP | 0.3 | 0.96 | 0.31 |
| sag[post]:Interf[high]:TDist[long]:aCP:nCP | -0.53 | 0.99 | -0.54 |
| lat[left]:Interf[high]:TDist[long]:aCP:nCP | -2.3 | 0.96 | -2.4 |
| lat[right]:Interf[high]:TDist[long]:aCP:nCP | -0.25 | 0.96 | -0.26 |
| sag[ant]:lat[left]:Interf[high]:TDist[long]:aCP:nCP | -0.63 | 1.4 | -0.46 |
| sag[post]:lat[left]:Interf[high]:TDist[long]:aCP:nCP | 1.2 | 1.4 | 0.88 |
| sag[ant]:lat[right]:Interf[high]:TDist[long]:aCP:nCP | 3.3 | 1.4 | 2.4 |
| sag[post]:lat[right]:Interf[high]:TDist[long]:aCP:nCP | -2.4 | 1.4 | -1.7 |

**Table S3:**
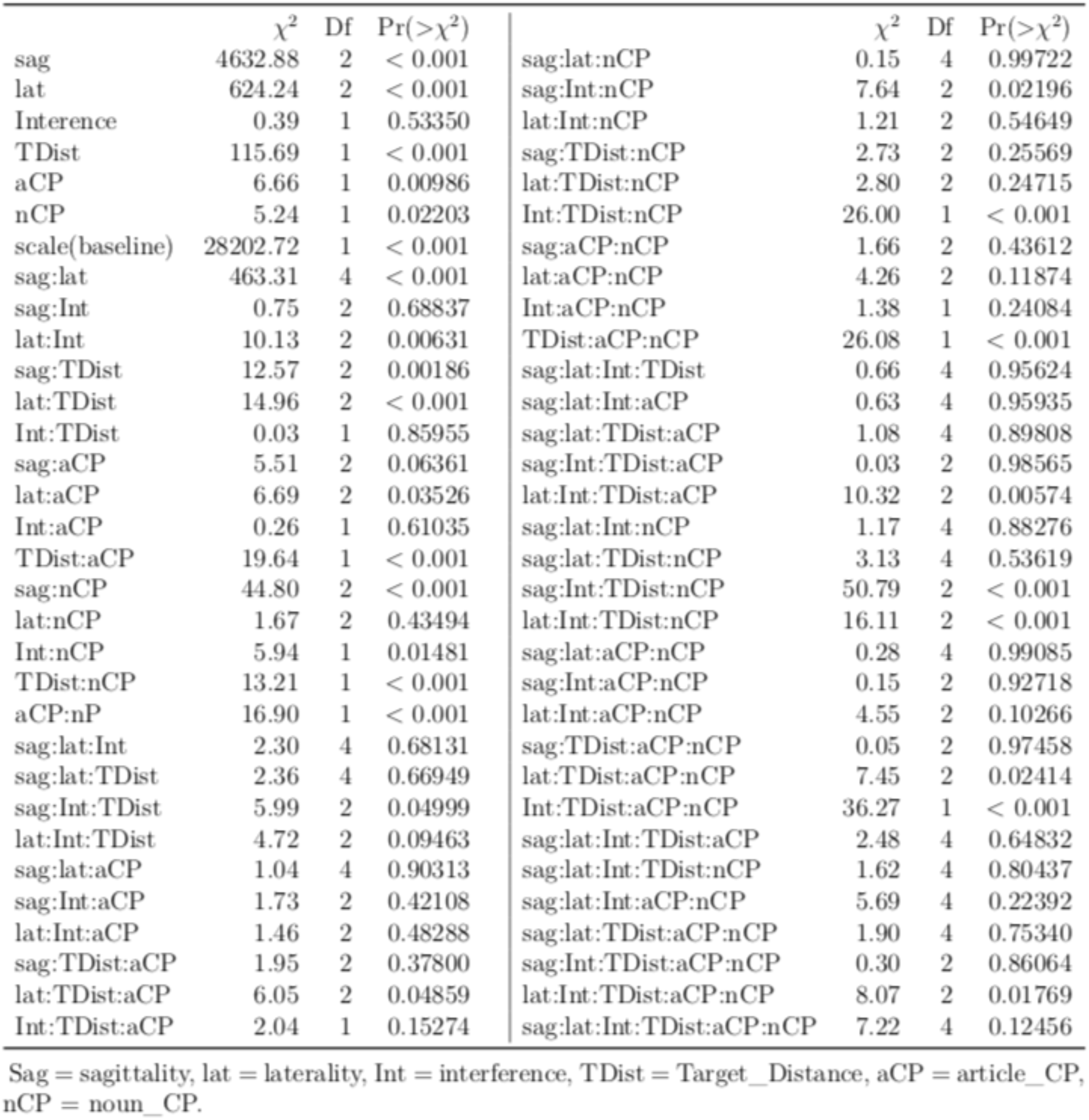
Summary of effects in the N400 time window after Noun onset (Type II Wald).

| | $\chi^2$ | Df | $\Pr(>\chi^2)$ | | $\chi^2$ | Df | $\Pr(>\chi^2)$ |
| --- | --- | --- | --- | --- | --- | --- | --- |
| sag | 4632.88 | 2 | < 0.001 | sag:lat:nCP | 0.15 | 4 | 0.99722 |
| lat | 624.24 | 2 | < 0.001 | sag:Int:nCP | 7.64 | 2 | 0.02196 |
| Interence | 0.39 | 1 | 0.53350 | lat:Int:nCP | 1.21 | 2 | 0.54649 |
| TDist | 115.69 | 1 | < 0.001 | sag:TDist:nCP | 2.73 | 2 | 0.25569 |
| aCP | 6.66 | 1 | 0.00986 | lat:TDist:nCP | 2.80 | 2 | 0.24715 |
| nCP | 5.24 | 1 | 0.02203 | Int:TDist:nCP | 26.00 | 1 | < 0.001 |
| scale(baseline) | 28202.72 | 1 | < 0.001 | sag:aCP:nCP | 1.66 | 2 | 0.43612 |
| sag:lat | 463.31 | 4 | < 0.001 | lat:aCP:nCP | 4.26 | 2 | 0.11874 |
| sag:Int | 0.75 | 2 | 0.68837 | Int:aCP:nCP | 1.38 | 1 | 0.24084 |
| lat:Int | 10.13 | 2 | 0.00631 | TDist:aCP:nCP | 26.08 | 1 | < 0.001 |
| sag:TDist | 12.57 | 2 | 0.00186 | sag:lat:Int:TDist | 0.66 | 4 | 0.95624 |
| lat:TDist | 14.96 | 2 | < 0.001 | sag:lat:Int:aCP | 0.63 | 4 | 0.95935 |
| Int:TDist | 0.03 | 1 | 0.85955 | sag:lat:TDist:aCP | 1.08 | 4 | 0.89808 |
| sag:aCP | 5.51 | 2 | 0.06361 | sag:Int:TDist:aCP | 0.03 | 2 | 0.98565 |
| lat:aCP | 6.69 | 2 | 0.03526 | lat:Int:TDist:aCP | 10.32 | 2 | 0.00574 |
| Int:aCP | 0.26 | 1 | 0.61035 | sag:lat:Int:nCP | 1.17 | 4 | 0.88276 |
| TDist:aCP | 19.64 | 1 | < 0.001 | sag:lat:TDist:nCP | 3.13 | 4 | 0.53619 |
| sag:nCP | 44.80 | 2 | < 0.001 | sag:Int:TDist:nCP | 50.79 | 2 | < 0.001 |
| lat:nCP | 1.67 | 2 | 0.43494 | lat:Int:TDist:nCP | 16.11 | 2 | < 0.001 |
| Int:nCP | 5.94 | 1 | 0.01481 | sag:lat:aCP:nCP | 0.28 | 4 | 0.99085 |
| TDist:nCP | 13.21 | 1 | < 0.001 | sag:Int:aCP:nCP | 0.15 | 2 | 0.92718 |
| aCP:nP | 16.90 | 1 | < 0.001 | lat:Int:aCP:nCP | 4.55 | 2 | 0.10266 |
| sag:lat:Int | 2.30 | 4 | 0.68131 | sag:TDist:aCP:nCP | 0.05 | 2 | 0.97458 |
| sag:lat:TDist | 2.36 | 4 | 0.66949 | lat:TDist:aCP:nCP | 7.45 | 2 | 0.02414 |
| sag:Int:TDist | 5.99 | 2 | 0.04999 | Int:TDist:aCP:nCP | 36.27 | 1 | < 0.001 |
| lat:Int:TDist | 4.72 | 2 | 0.09463 | sag:lat:Int:TDist:aCP | 2.48 | 4 | 0.64832 |
| sag:lat:aCP | 1.04 | 4 | 0.90313 | sag:lat:Int:TDist:nCP | 1.62 | 4 | 0.80437 |
| sag:Int:aCP | 1.73 | 2 | 0.42108 | sag:lat:Int:aCP:nCP | 5.69 | 4 | 0.22392 |
| lat:Int:aCP | 1.46 | 2 | 0.48288 | sag:lat:TDist:aCP:nCP | 1.90 | 4 | 0.75340 |
| sag:TDist:aCP | 1.95 | 2 | 0.37800 | sag:Int:TDist:aCP:nCP | 0.30 | 2 | 0.86064 |
| lat:TDist:aCP | 6.05 | 2 | 0.04859 | lat:Int:TDist:aCP:nCP | 8.07 | 2 | 0.01769 |
| Int:TDist:aCP | 2.04 | 1 | 0.15274 | sag:lat:Int:TDist:aCP:nCP | 7.22 | 4 | 0.12456 |

Sag = sagittality, lat = laterality, Int = interference, TDist = Target\_Distance, aCP = article\_CP, nCP = noun\_CP.

**Table S4:**
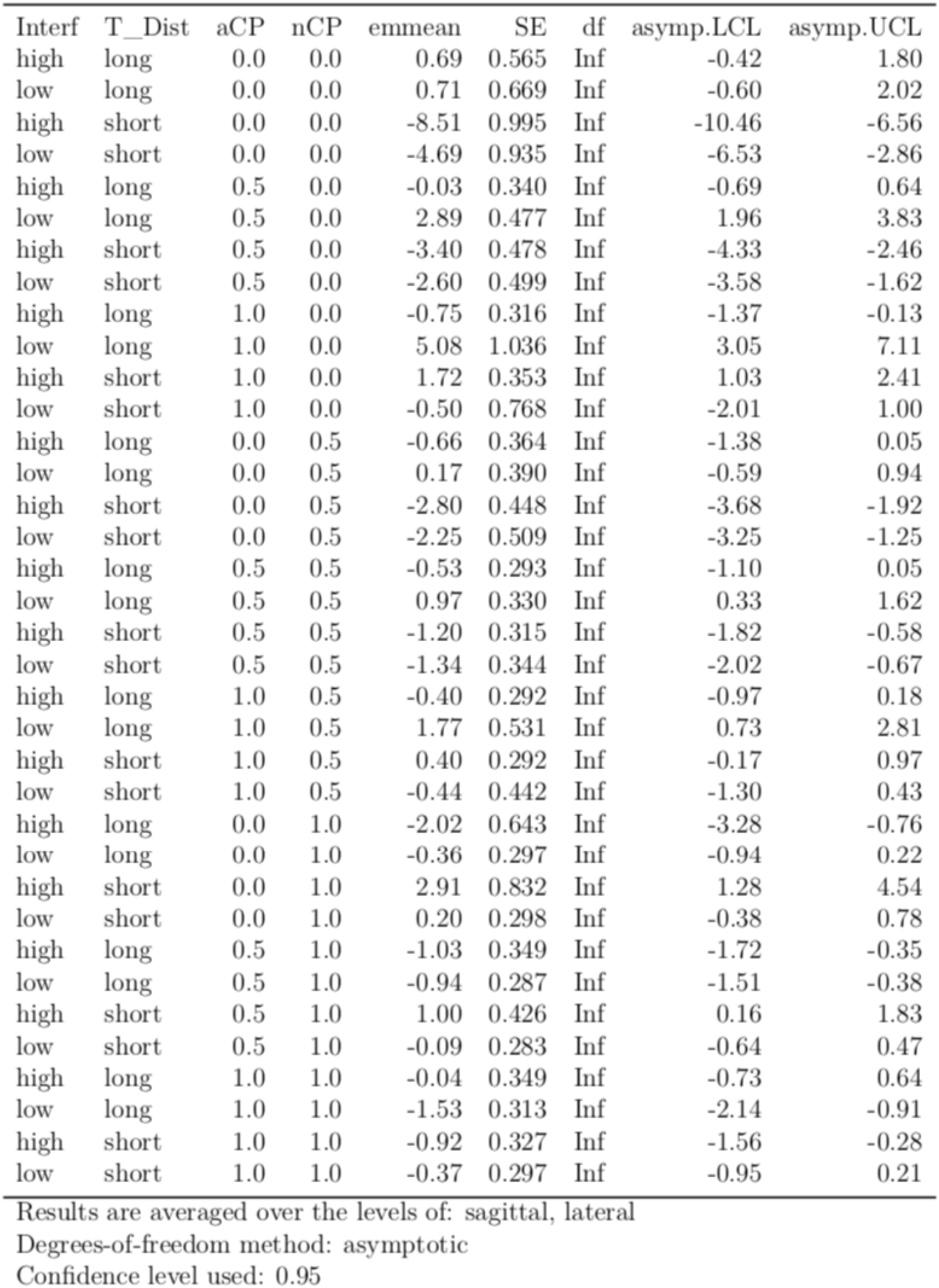
Estimated marginal means for the Noon in the N400 time window produced by the call `emmeans(noun_n4_Ionly, c(“Interference”, “Target_Distance”, “article_CP”, “nounCP”), at = list(article_CP = c(0, 0.5, 1), article_CP = c(0, 0.5, 1)))`. Interf = Interference, T_Dist = Target_Distance, aCP = article_CP. nCP= nonn_CP.

**Figure S2:**
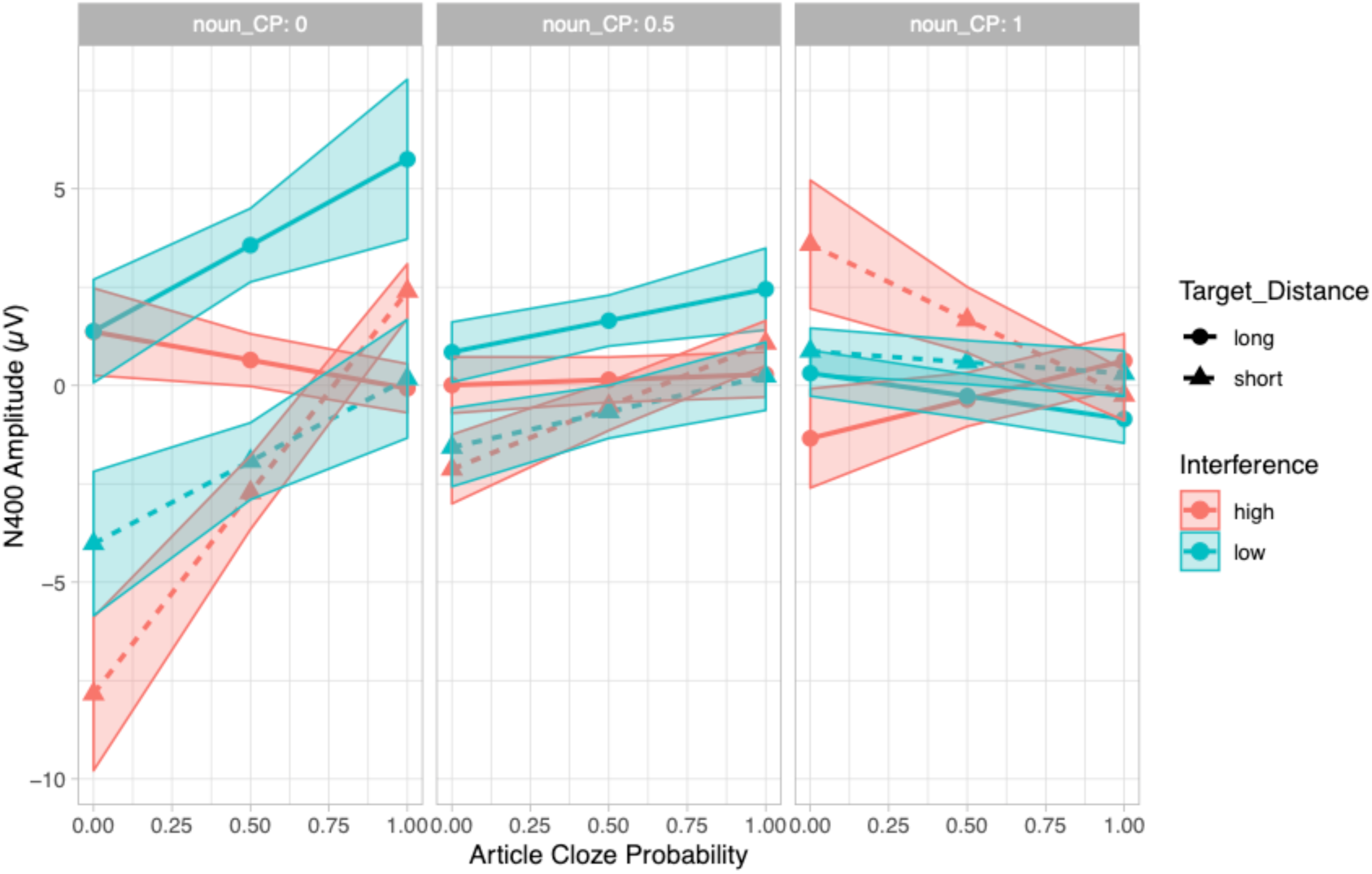
Predicted values of mean amplitude in the N400 time window at noun position by interference, target distance, article CP and noun CP. Shaded regions indicate 95% confidence intervals.

When both Article CP and Noun CP are low, the short target distance conditions are more negative than the long target distance conditions – irrespective of interference. When Article CP is high and Noun CP is low, the high interference conditions and the low interference / short target distance condition are more negative than the low interference / long target distance condition. As Noun CP increases, the interference and target distance effects decrease depending on Article CP. When Noun CP is high and Article CP is low, the high interference / long target distance condition and the two low interference conditions are more negative than the high interference / short target distance condition. When both Noun CP and Article CP are high, no effects were apparent.

It was mentioned in the Methods section that the combinations of CP values are not evenly distributed across conditions; therefore we present the data basis for the above-described results in Figure S2. Similar to the article effects and their data basis, for the high interference conditions at the noun, Article CP ranged from approx. 0.5 to 1 across the Noun CP scale. Additionally, for the low interference conditions, Noun CP ranged from approx. 0.5 to 1.

**Figure S2:**
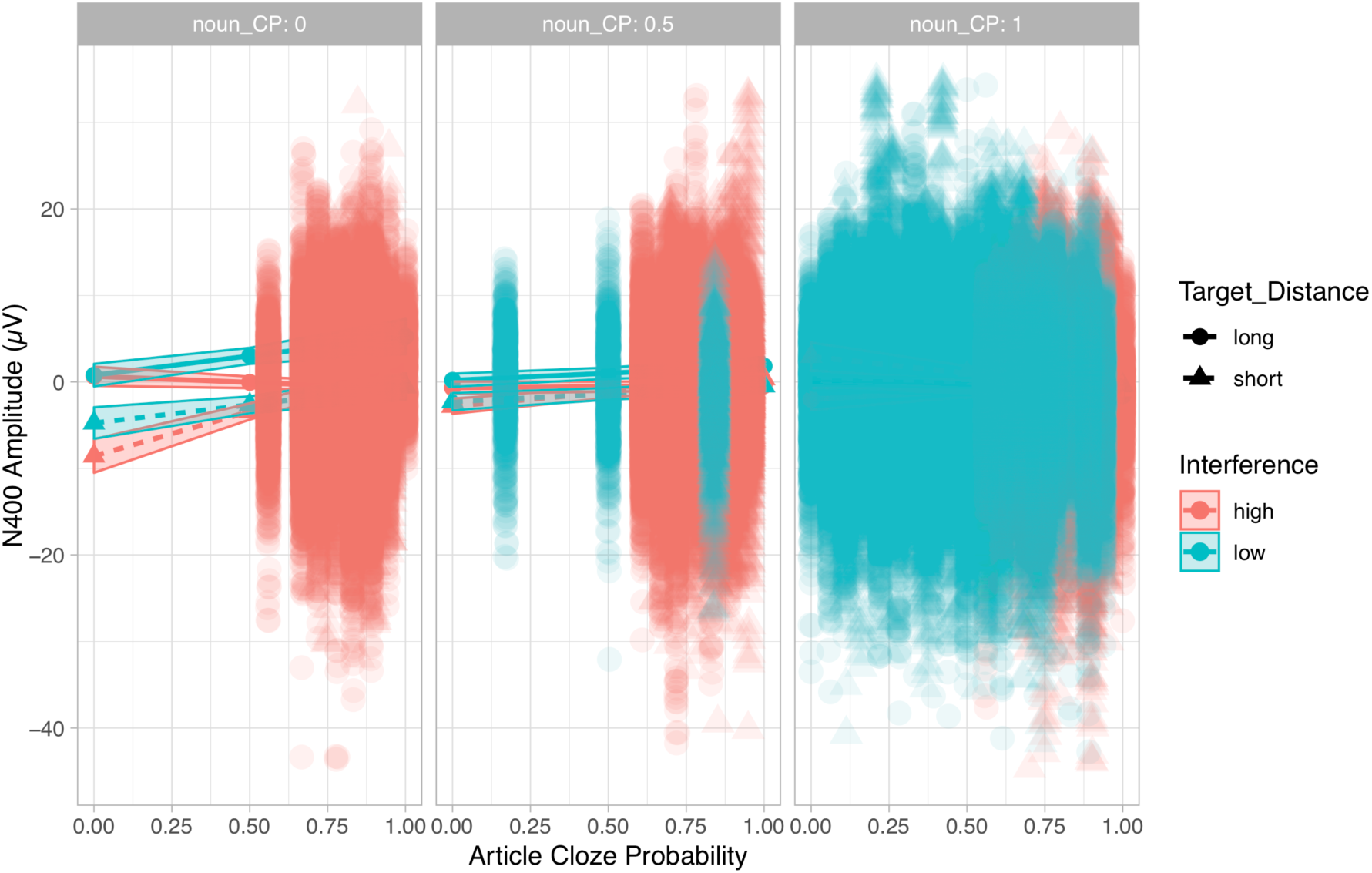
Distribution of data points with estimated effects at noun position in the background.

